# Investigation on How Dynamic Effective Connectivity Patterns Encode the Fluctuating Pain Intensity in Chronic Migraine

**DOI:** 10.1101/2022.02.23.481583

**Authors:** Iege Bassez, Frederik Van de Steen, Sophie Hackl, Pauline Jahn, Astrid Mayr, Daniele Marinazzo, Enrico Schulz

## Abstract

Chronic migraine is characterised by persistent headaches for more than 15 days per month; the intensity of the pain is fluctuating over time. Here, we explored the dynamic interplay of connectivity patterns between regions known to be related to pain processing and their relation to the ongoing dynamic pain experience. We recorded EEG from 80 sessions (20 chronic migraine patients in 4 separate sessions of 25 minutes). The patients were asked to continuously rate the intensity of their endogenous headache. On different time-windows, a dynamic causal model (DCM) of cross spectral responses was inverted to estimate connectivity strengths. For each patient and session, the evolving dynamics of effective connectivity were related to pain intensities and to pain intensity changes by using a Bayesian linear model. Hierarchical Bayesian modelling was further used to examine which connectivity-pain relations are consistent across sessions and across patients.

The results reflect the multi-facetted clinical picture of the disease. Across all sessions, each patient with chronic migraine exhibited a distinct pattern of pain intensity-related cortical connectivity. The diversity of the individual findings are accompanied by inconsistent relations between the connectivity parameters and pain intensity or pain intensity changes at group level. This suggests a rejection of the idea of a common neuronal core problem for chronic migraine.

## 1 Introduction

The experience of pain is associated with a number of different neuronal oscillations in the brain (Michail et al., 2016; Ploner et al., 2017), but precisely how brain regions in chronic pain conditions are synchronised, and at which frequencies, remains largely unknown.

Across functional neuroimaging studies, several regions have been consistently found to be related to pain processing, i.e. the thalamus, the primary (S1) and secondary somatosensory cortices (S2), the insular cortex (IC), the anterior cingulate cortex (ACC), and the prefrontal cortex (PFC; Apkarian et al., 2005; Bushnell et al., 2013; Price, 2000; Rainville, 2002; Tracey, 2008; Treede et al., 1999). These brain regions are anatomically connected, allowing serial and parallel information transfer (Price, 2000). For the serial pathway, input relayed in the thalamus flows from S1 to S2, which plays a role in encoding sensory information of nociceptive input (Bushnell et al., 2013). This ascending pain pathway then continues from S2 to the IC, then to the ACC. In parallel, the S2, IC and ACC can also directly receive input from the thalamus (Frot et al., 2008; Liang et al., 2011).

The IC has an integrative function in pain processing (Brooks & Tracey, 2007); the anterior insular cortex (aIC) is functionally connected to regions related to affective and cognitive aspects of pain, and the posterior insular cortex (pIC) is predominantly connected to regions related to sensory aspects of pain (Peltz et al., 2011). The ACC is involved in encoding emotional and motivational aspects of pain and has bi-directional connections with the PFC (Bushnell et al., 2013; Price, 2000; Rainville, 2002; Wiech, Ploner, et al., 2008).

The medial and lateral PFC have been functionally differentiated; medial PFC (mPFC) activity has been positively associated with pain intensity (Baliki et al., 2006; Hashmi et al., 2013; May et al., 2019; Nickel et al., 2017; Schulz et al., 2015), and dorsolateral PFC (DLPFC) activity has been negatively associated with pain affect, where it plays an important role in pain modulation (Lorenz et al., 2003; Wiech, Farias, et al., 2008). Several other studies utilising effective connectivity methods have found that placebo analgesia (i.e., endogenous pain modulation) increased connectivity from the DLPFC to the dorsal ACC (dACC; Craggs et al., 2007; Sevel et al., 2015). Similarly, when investigating connectivity between the left and right DLPFC, Sevel and colleagues found that the higher the connectivity strength from the right to left DLPFC, the higher the temperatures of the stimuli were in order to be perceived as painful (Sevel et al., 2016). This suggests an important role of intra- and interhemispheric connectivity in pain modulation.

Importantly, structural and functional abnormalities have been reported in these pain-related areas in patients with migraine (Borsook et al., 2016; Filippi & Messina, 2019; Jia & Yu, 2017; Tolner et al., 2019). Using a resting-state design, Lee and colleagues found that compared to episodic migraine patients, chronic migraine (CM) patients have stronger connectivity in brain regions that included the ACC, aIC and DLPFC (Lee et al., 2019). In these regions, grey matter reductions have also been shown in migraine patients (Kim et al., 2008; Maleki et al., 2012; Rocca et al., 2006). Although referring to these regions as pain-related regions, they are certainly not pain specific, instead the perception of pain likely arises from interactions between brain regions, resulting in a distinct spatial pattern of neural activity (Kucyi & Davis, 2015; Liang et al., 2019).

In order to gain insight into how the interactions between these regions encode the pain experienced by migraine patients, we investigated how dynamic effective connectivity between pain-related regions relates to the ongoing and fluctuating headache in CM. We hypothesised that connection strengths in the ascending pain pathways would be enhanced with increasing pain or higher levels of pain (i.e., S1 > S2, S2 > S1, S2 > pIC, pIC > aIC, aIC > dACC, pIC > dACC, dACC > mPFC, mPFC > dACC, dACC > DLPFC in both hemispheres). In contrast, we expected connection strengths from regions involved in pain modulation to be enhanced with decreasing pain or lower levels of pain (i.e., DLPFC > dACC in both hemispheres, left DLPFC > right DLPFC, right DLPFC > left DLPFC). Finally, we aimed to give a more detailed insight into the connectivity effects in terms of spectral outcomes in each region.

## 2 Methods

### 2.1 Participants

Twenty CM patients (18 females, aged 34±13 years) participated in this study. All participants gave written informed consent. The study was approved by the Ethics Committee of the Medical Department of the Ludwig-Maximilians-Universität München and conducted in conformity with the Declaration of Helsinki. The patients were diagnosed according to the ICHD-3 (Headache Classification Committee of the International Headache Society (IHS), 2018), defined as a headache occurring on 15 or more days/month for more than 3 months, which, on at least 8 days/month, has the features of migraine headache (mean CM: 15±12 years). The patients in this study had a history of migraine attacks between 2 and 50 years (*M* = 15.10 years, *SD* = 12.01 years). The mean pain intensity as specified in the questionnaires was 4.90 (*SD* = 1.30) on a scale to 10. All patients were seen in a tertiary headache centre.

All patients were permitted to continue their pharmacological treatment at a stable dose (Supplementary Table 1). The patients did not report any other neurological or psychiatric disorders or had contraindications for an MRI examination. Patients who had any additional pain were excluded. For all patients, the pain was fluctuating and not constant at the same intensity level. Patients with no pain or migraine attacks on the day of the measurement were asked to return on a different day. Patients were characterised using the German Pain Questionnaire (Deutscher Schmerzfragebogen; Casser et al., 2012) and the German version of the Pain Catastrophizing Scale (PCS; Supplementary Table 1; Sullivan et al., 1995). The pain intensity describes the average pain in the last 4 weeks from 0 to 10 with 0 representing no pain and 10 indicating maximum imaginable pain (please note that this scale differs from the one used in the EEG experiment). The German version of the Depression, Anxiety and Stress Scale (DASS) was used to rate depressive, anxiety, and stress symptoms over the past week (Lovibond & Lovibond, 1995). None of the patients was excluded based on their questionnaire scores. A study on healthy subjects found similar results: a large sample of 1794 participants reported scores for depression of 3 ± 4, for anxiety of 2 ± 3, and for stress of 5 ± 4 (Henry & Crawford, 2005). None of the patients in our study reported any psychiatric comorbidity. Patients were compensated with 60€ for each session. The patients were recorded four times across 6 weeks with a gap of at least 2 days (12±19 days) between sessions.

### 2.2 Experimental procedure

During each EEG recording session, patients continuously rated the intensity of their ongoing headache for a duration of 25 minutes using a linear slider potentiometer (Jahn et al., 2021; Mayr et al., 2021, 2022; Schulz et al., 2020). The pain scale ranged from 0 to 100 with ‘0’ representing no pain and ‘100’ representing highest experienced pain. The patients could see their ratings on the monitor as the position of the red cursor on a horizontal grey bar (visual analogue scale) and numerically (in steps of 5; numeric analogue scale). The patients were instructed to rate their pain as quickly and accurately as possible, to move as little as possible during the recording, and to focus on their varying pain. The pain ratings of the 20 CM patients across the four sessions are plotted in Supplementary Figure 1.

### 2.3 Data acquisition

The EEG data were recorded using an array of 64 equidistantly distributed electrodes (EASYCAP, Brain Products GmbH, Germany). The EEG was referenced to a vertex electrode, grounded at the nose and sampled at 1 kHz. Impedances were kept below 20 kΩ. Individual electrode positions were acquired using a stereo-optical system (CapTrak, Brain Products GmbH, Germany). During one of the visits, structural MRI (T1-weighted) images were acquired with a 3 tesla MRI scanner (Magnetom Skyra, Siemens, Germany) using a 64-channel head coil. The following parameters were used: TR/TE = 2060/2.17 ms; flip angle = 12°; number of slices = 256; slice thickness = 0.75 mm; FoV = 240 × 240.

### 2.4 Pre-processing

The raw EEG data were pre-processed in the BrainVision Analyzer software (Brain Products GmbH, Germany). Bad channels were interpolated, data were high-pass filtered with a lower cut-off at 1 Hz and decomposed using an independent component analysis (ICA). On the component data, 50 Hz power line noise was removed using CleanLine (Mullen, 2012) and artefactual components reflecting eye movements and other larger artefacts were removed from the data. A second ICA was performed and muscle artefacts were removed from the data (Liebisch et al., 2021). A third ICA was utilised to remove components with residual artefacts and spectrum interpolation (Leske & Dalal, 2019) eliminated residual power line noise. Finally, the data were downsampled to 256 Hz and re-referenced to the common average reference.

### 2.5 Effective connectivity and pain

In order to investigate how fluctuations in effective connectivity are related to the patients’ endogenous dynamic pain experience, we used dynamic causal modelling (DCM) and a hierarchical Bayesian framework as described and used previously (Park et al., 2018; Van de Steen et al., 2019). More specifically, we used DCM for cross spectral density (CSD) data features (K. J. Friston et al., 2012) combined with multilevel parametric empirical Bayes (PEB; K. J. Friston et al., 2016).

#### 2.5.1 Within-window level: DCM for cross spectral density data features

The preprocessed data were imported to SPM12 (Penny et al., 2011) running on MATLAB (Mathworks, USA; version 2017b). DCM for CSD was performed on every 5 second EEG window. The preprocessed time-series were transformed to cross-spectral densities between 4 to 90 Hz using a vector autoregressive model of order 12. DCM for CSD aims to explain how the CSD data features are generated by underlying neurophysiology by using a biologically plausible generative model (neural model/state equations + forward model/observation equations). Regions and connections between them must be defined *a priori* in the model. By inverting the generative model using a variational Bayesian optimisation scheme (Variational Laplace algorithm; K. Friston et al., 2007a), we derived the posterior distribution of the connectivity parameters defined in the model. The optimisation scheme uses free energy as the objective function, which approximates the log model evidence (K. Friston et al., 2007b).

We used a convolution-based neural mass model where each source (i.e., a functionally specialised brain region) has three neuronal subpopulations: excitatory spiny stellate cells, inhibitory interneurons, and excitatory pyramidal cells (Moran et al., 2013). Coupling between sources can be divided into forward, backward and lateral extrinsic connections, based on what the seed and target neuronal subpopulations are. The extrinsic connection types to be defined in the model can be established based on the hierarchical organisation of the cortex (Felleman & Van Essen, 1991).

For the spatial forward model, each source was treated as a patch on the cortical surface (`IMG’ option in SPM12) with a radius of 10 mm (Daunizeau et al., 2009). Individual structural MRI (T1-weighted) images were used to compute individual cortical meshes and co-registration was performed with individual electrode positions. Volume conduction models of the head were constructed based on the boundary element method (BEM). For three patients, we were unable to record individual electrode positions and thus standard coordinates were used. For every patient, the individual head model with the different tissue types (brain, skull and scalp), the normalised individual cortical mesh, and individual electrode locations were plotted after co-registration for verification. Default prior parameters for the generative model were used.

The selection of pain-related regions and their connectivity pattern was based on previous literature (Apkarian et al., 2005; Bushnell et al., 2013; Craggs et al., 2007; Lorenz et al., 2003; Price, 2000; Rainville, 2002; Schulz et al., 2015; Sevel et al., 2015, 2016; Tracey, 2008; Treede et al., 1999; Wiech, Farias, et al., 2008). The network that was examined included the left and right primary somatosensory cortex (S1), left and right secondary somatosensory cortex (S2), left and right anterior and posterior insular cortex (aIC; pIC), dorsal anterior cingulate cortex (dACC), medial prefrontal cortex (mPFC), and the left and right dorsolateral prefrontal cortex (DLPFC). Coordinates for the left and right DLPFC were based on Sevel et al. and coordinates for the mPFC were based on Schulz et al. (Schulz et al., 2015; Sevel et al., 2015). For the other regions, centre coordinates were determined by termed-based meta-analyses in Neurosynth (Neurosynth.org; Yarkoni et al., 2011). All coordinates were verified by atlases in FSL. For the dACC and mPFC, we treated the left and right cortex as a single source; given their proximity and the spatial resolution of EEG, they are difficult to separate. In Supplementary Table 2, the centre MNI coordinates of the regions are given and in Figure 1, the presumed coupling between the regions is illustrated.

**Figure 1.**
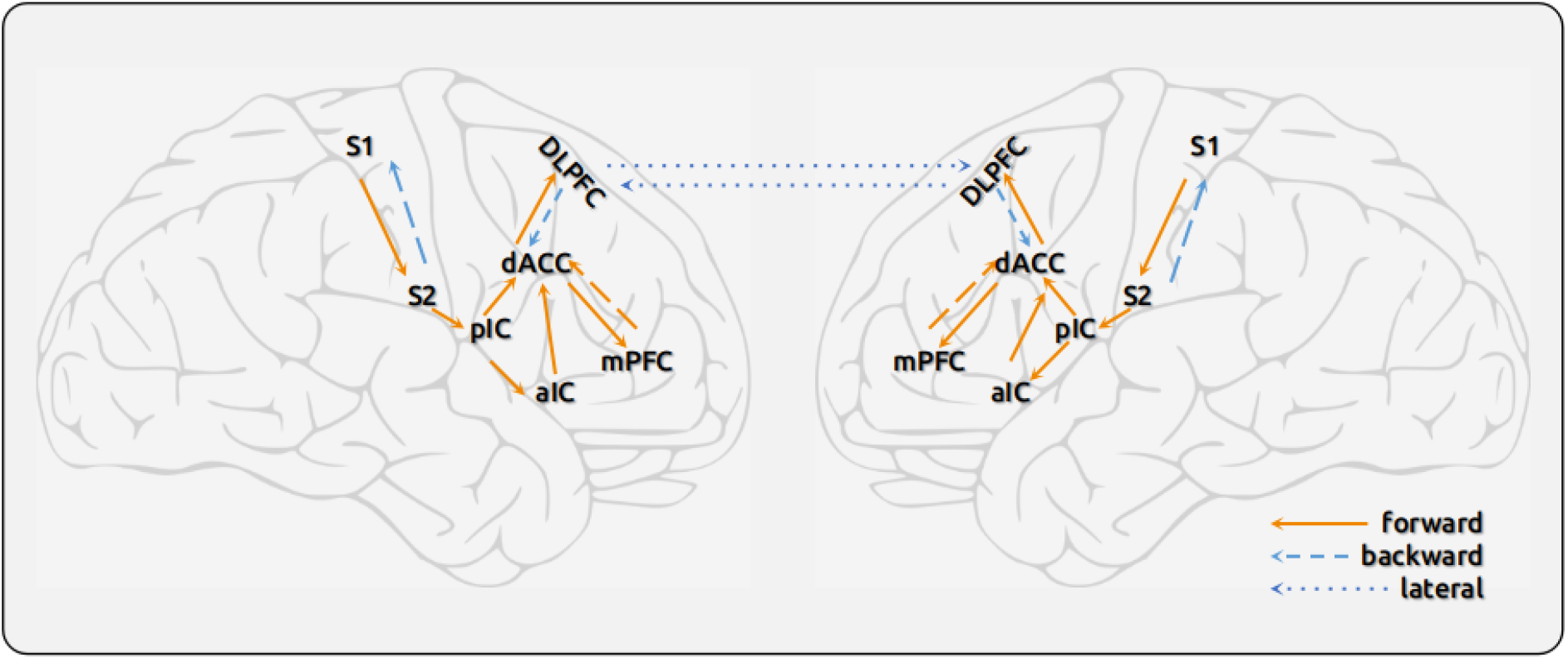
Forward, backward and lateral connections that were estimated in each time window. S1, primary somatosensory cortex; S2, secondary somatosensory cortex; pIC, posterior insular cortex, aIC, anterior insular cortex; dACC, dorsal anterior cingulate cortex; mPFC, medial prefrontal cortex; DLPFC, dorsolateral prefrontal cortex.

At this level, the specified model was inverted for each time-window independently in order to estimate effective connectivity between selected regions (see Figure 1) in a certain pain state. To invert the DCM models, the function ‘spm_dcm_csd’ was used. The actual inversions of the DCMs were performed in parallel on the high-performance computing infrastructure of Ghent University, the Free University of Brussels, and on the Linux cluster of the Leibniz Supercomputing Centre. The same version of SPM (r7771) was used on all clusters but the MATLAB version differed (2017b, 2019a). Running the same window on the different clusters resulted in similar results. The explained variance of each model was calculated to investigate whether the DCM was able to fit the observed CSD well. Overall, we treated models with an explained variance of 50% and more as adequate for further analysis. If the explained variance did not reach 50%, the model was excluded from further analysis.

#### 2.5.2 Within-session level: PEB across windows

Here, we estimated the connectivity-pain rating relations for each session of each participant separately. The rating data were continuously recorded with a variable sampling rate. As a DCM was inverted on every 5 second window, we processed the rating data accordingly. The ratings within a 5 second window were averaged to have an estimate of the pain state during that time window.

To disentangle the distinct aspects of pain intensity (AMP - amplitude) from cortical processes related to the sensing of rising and falling pain, we computed the ongoing rate of change in the pain ratings. This vector is calculated as the slope of the regression of the least-squares line across a 5 s time window (SLP - slope, encoded as 1, -1, and 0). Periods coded as 0 indicate time frames of constant pain. The absolute slope of pain ratings (aSLP - absolute slope, encoded as 0 and 1) represents periods of motor-related connectivity (slider movement), changes of visual input (each slider movement changes the screen), and decision-making (each slider movement prerequisites a decision to move). Periods coded as 0 indicate time frames of constant pain without the need to move the slider. See Supplementary Figure 1 for the detailed rating time courses of each session for each subject.

Moving the slider to indicate a change in pain is embedded in a cascade of interwoven steps. Preceding functional connections can influence the current rating and the current rating can have an impact on subsequent cortical connectivity. To account for the unknown timing between brain dynamics and the subsequent ratings, we shifted the rating vectors (AMP, SLP, and aSLP) between -15 and 20 seconds in steps of 1 second. Therefore, each statistical model was computed 36 times along the time shifts of the rating vector.

We modelled the connectivity parameters over time-windows using Bayesian linear models where the endogenous pain rating vectors (AMP, SLP, and aSLP) were used as regressors. In total, 36 models (shifts from -15 to 20 seconds in steps of 1 second) were estimated for each session of each participant. The structure of each Bayesian linear model is described below:

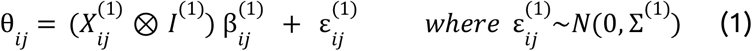

In equation (1), *θ*_*ij*_ are the vectorised DCM parameters (i.e., estimates of all connection strengths) for session *i* of participant *j*. Note that in this model the uncertainties of the estimated connection strengths (i.e., posterior covariance) are considered. The parameters are organised in such a way that the first elements of *θ*_*ij*_ are the DCM parameters of the first window and the following elements are the DCM parameters of the second window and so on. The size of *θ*_*ij*_ thus, equals the number of parameters (P) times the number of windows (W) in the session. 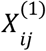 is the design matrix with W rows and 4 columns, where the first column is a column of ones, the second column is the AMP vector (standardised), the third column is the SLP vector, and the last column is the aSLP vector. *I*^(1)^ is the identity matrix of size P. The Kronecker tensor product (⊗) of 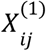 and *I*^(1)^ means that each connectivity parameter can show a relation with pain ratings. The last term, 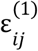 represents the between-window variability (random effects). The 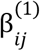 parameters (4P x 1) were estimated with PEB (K. J. Friston et al., 2016) using the ‘spm_dcm_peb’ function.

In the next step, we compared the different shift models. We averaged the free energies of the models over sessions and participants and compared each model with the worst model (approximate log Bayes factor). The winning model was further used.

#### 2.5.3 Within-subject level: PEB across sessions

Here, we investigated for each participant which connectivity-pain rating relations (from the winning shift model) are consistent over sessions (non-trivial across session means) using the following PEB model:

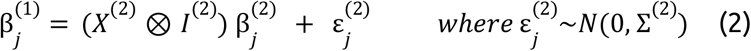

In equation (2), 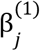 are the vectorised connectivity-pain rating relations for all four sessions estimated at the previous level for participant *j*. 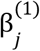 thus equals the number of connectivity parameters in the network times the number of regressors in the previous level (4P) times the number of sessions (S). *X*^(2)^ is the design matrix with 4 rows and 1 column. *I*^(2)^ is the identity matrix of size P. The last term, 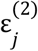 represents the between-session variability (random effects). The average connectivity-pain rating relations (i.e., 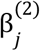 parameters of size 4P) were estimated using the ‘spm_dcm_peb’ function.

#### 2.5.4 Within-group level: PEB across subjects

Here, we investigated which connectivity-pain rating relations are systematic over patients. Therefore, participants’ results that were consistent across sessions were entered in a group-level Bayesian linear model:

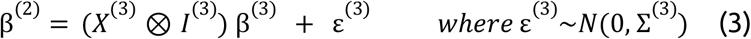

In equation (3), β^(2)^ are the vectorised parameter estimates. The size of β^(2)^ equals the number of parameters in the network times the regressors of previous levels (4P) times the number of participants (N). The group design matrix *X*^(3)^with N rows, consists of a column with constants to model the average of the pain-related changes in effective connectivity across participants.*I*^(3)^ is the identity matrix of size 4P. ε^(3)^represents the between-subject variability (random effects).

Bayesian model reduction (BMR) and a greedy search were used in order to remove parameters that are redundant in the full model (K. J. Friston et al., 2016; K. Friston & Penny, 2011). The functions ‘spm_dcm_peb’ and ‘spm_dcm_peb_bmc’ were used.

#### 2.5.5 Sensitivity analysis

We performed a sensitivity analysis that shows the change in outcome measures as a function of a change in parameters values. Technically, we numerically evaluated (at the posterior mean) the gradient of outcomes with respect to the connectivity parameters that show a pain-related connectivity at the group level (i.e., posterior probability > .95). For simplicity, our outcome measures were the simulated spectral density of a “virtual local field potential” in each region. This was conducted for every DCM separately. Sensitivity profiles within participants were averaged. As a result, we obtained a sensitivity profile per participant (size: number of frequency bands by number of regions by number of parameters). Permutation one sample t-tests combined with the maximum statistic approach to correct for multiple testing, were performed to assess significant sensitivity profiles at the group-level (Maris & Oostenveld, 2007). This analysis was included to give us a more detailed insight into the connectivity effects in terms of spectral outcomes in each source.

## 3 Results

### 3.1 DCM results

Across participants, the average number of remaining windows for session 1 to 4 were 287.10 (*SD* = 14.82), 285.15 (*SD* = 19.12), 287.55 (*SD* = 15.20) and 285.80 (*SD* = 26.99), respectively. The mean percentages explained variance of these windows for session 1 to 4 were 98.10 (*SD* = 0.98), 98.02 (*SD* = 0.99), 98.10 (*SD* = 0.92), and 97.96 (*SD* = 1.17), respectively. The number of remaining windows for each session of each patient and the average explained variance over these windows for each session are given in Supplementary Table 2.

### 3.2 PEB results

#### 3.2.1 Group results

Comparison of the 36 shift models revealed the 0 second shifting as the winning model. In Figure 2, the log Bayes factors (free energy model *i* minus free energy of the worst model) are plotted. The model with the lowest free energy was the model with the largest negative shift (−15 sec).

**Figure 2.**
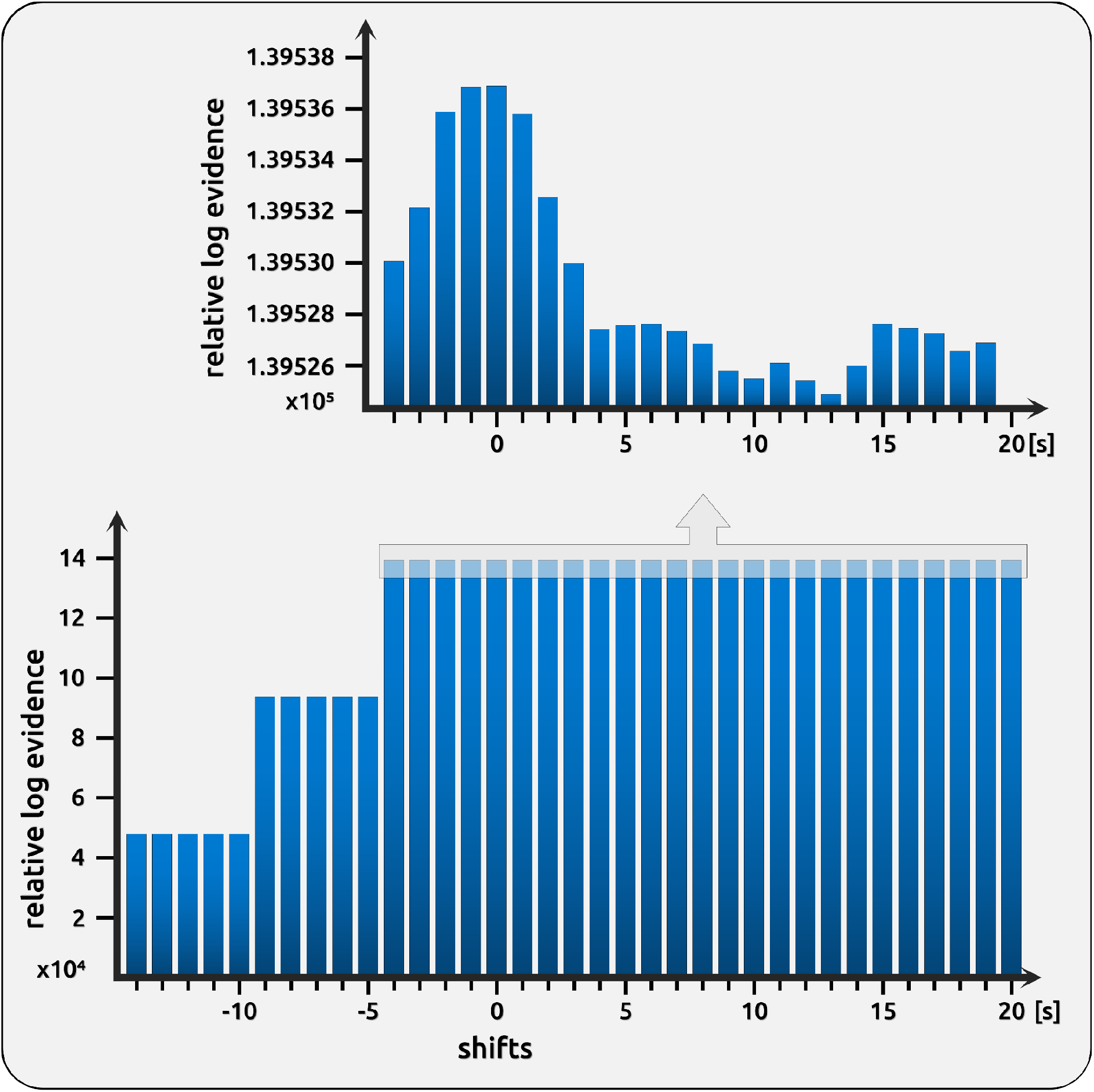
Bayesian model comparison of the different shift models from -15 to 20 seconds. The log Bayes factors (free energy model *i* minus free energy of the worst model) are plotted. The model with the lowest free energy was the model with the largest negative shift (−15 sec).

PEB results before and after BMR showed no consistent relations between the connectivity parameters and pain intensities at group level (all posterior probabilities of being different from zero <= 0.95). One connection of note is the connectivity from the DLPFC-r to the DLPFC-l, which showed on average a negative relation with AMP (β = -0.060, posterior probability = 0.92), but was pruned away after BMR. Connectivity changes related to increasing and decreasing pain were not consistent at group level (all posterior probabilities < 0.95). Likewise, one connection from the dACC to DLPFC-l showed a negative relation with increasing pain (β= -0.192, posterior probability = 0.95). This connection was pruned away after BMR. Some connectivity parameters were related to aSLP. Before BMR, the following connections had negative relations (posterior probability > 0.95) with aSLP: the connection from S1-l to S2-l (β= -0.277, posterior *SD* β = 0.132), from S2-l to pIC-l (β= -0.279, posterior *SD* β = 0.127), from aIC-l to dACC (β= -0.244, posterior *SD* β = 0.126) and from pIC-l to dACC (β= -0.251, posterior *SD* β = 0.124). The first three connections were still present after BMR.

#### 3.2.2 Individual results

In Figures 3 and 4 and Supplementary Figure 2, the individual results from the within-session level PEB and within-subject level PEB are displayed. For visual purposes, columns 1 to 4 show the beta values from the within-session level PEB and are thresholded at a posterior probability of higher than 0.75. Column 5 shows the beta values from the within-subject PEB, thresholded at a posterior probability higher than 0.95. The connectivity from the DLPFC-r to the DLPFC-l showed at group level a negative relation with AMP, but was pruned away after BMR. Looking at individual results (Figure 3), we see that two patients had this negative relation consistently across sessions whereas one patient showed an opposite relation. Other patients showed both negative and positive relations in some single sessions. On average, the connection from the dACC to the DLPFC-l showed a negative relation with increasing pain, however this connection was removed after BMR. Three patients showed this negative relation consistently across sessions, and one patient showed a positive relation across sessions. Other patients showed both negative and positive relations in single sessions (Figure 4). Supplementary Figure 2, shows the relations with aSLP.

**Figure 3.**
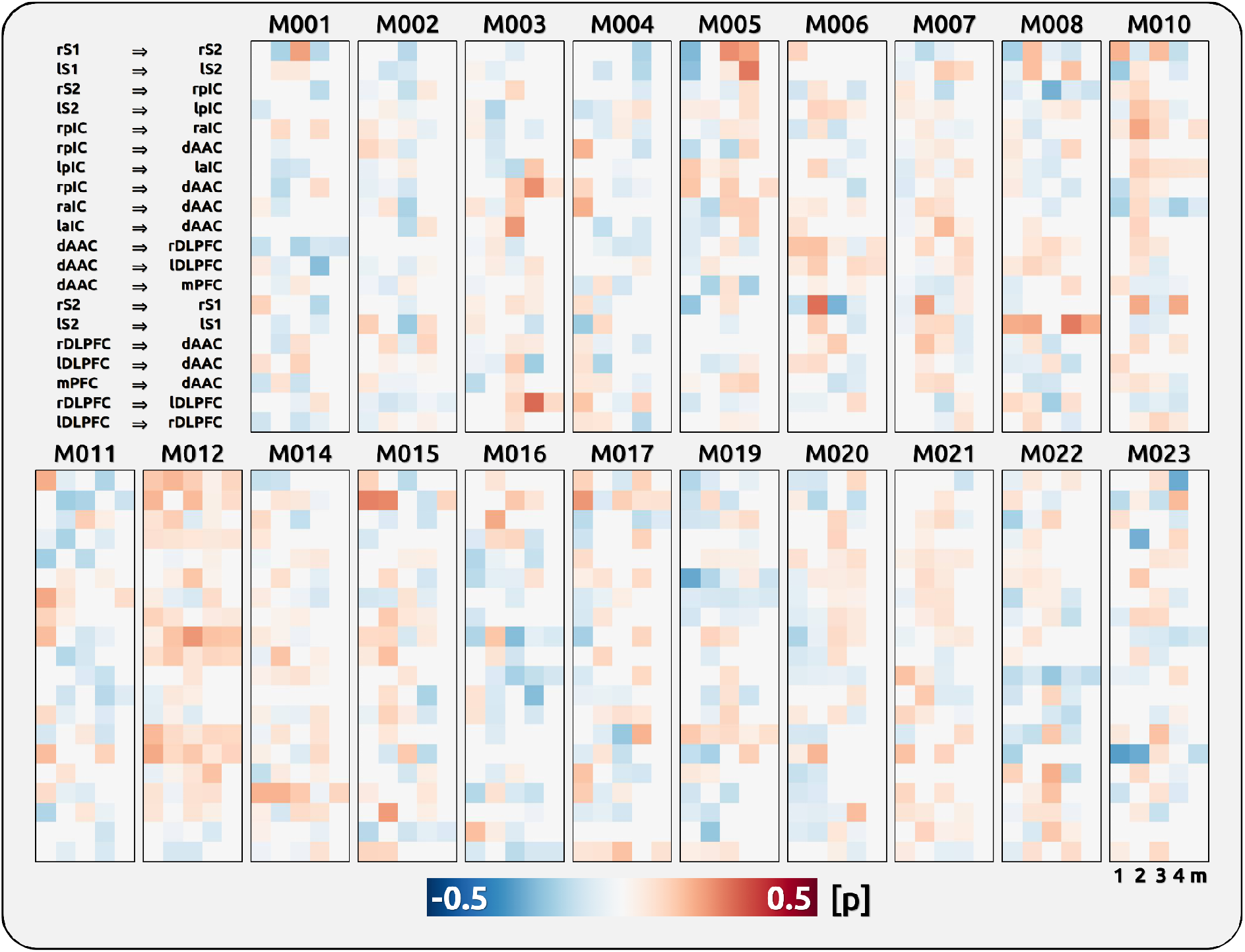
Individual results of relations between the connectivity parameters and AMP in each session and across sessions. Columns 1 to 4 represent the beta values from the within-session level PEB and are thresholded at a posterior probability > 0.75. Column 5 represents the beta values from the within-subject PEB, thresholded at a posterior probability > 0.95.

**Figure 4.**
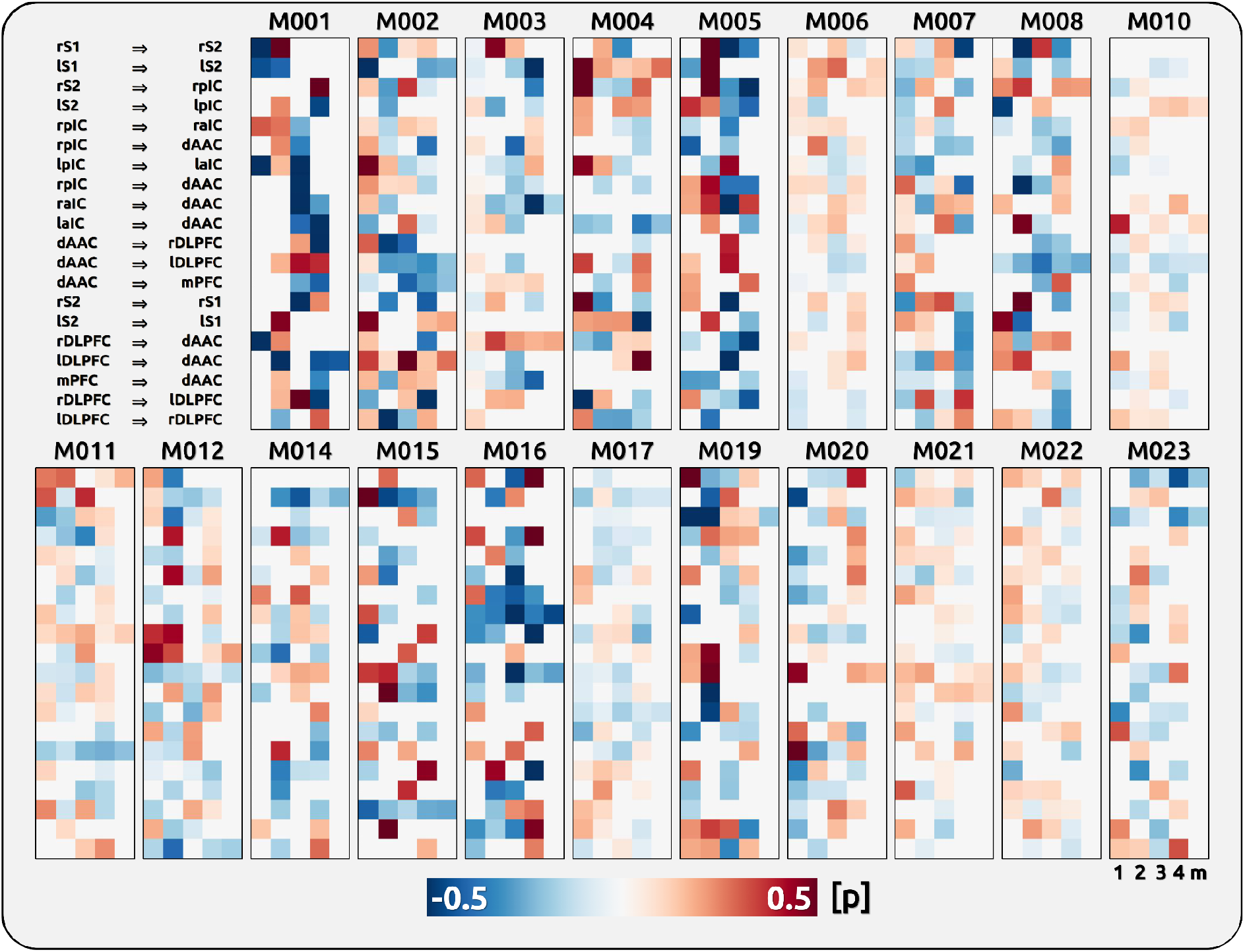
Individual results of relations between the connectivity parameters and SLP in each session and across sessions. Columns 1 to 4 represent the beta values from the within-session level PEB and are thresholded at a posterior probability > 0.75. Column 5 represents the beta values from the within-subject PEB, thresholded at a posterior probability > 0.95.

Although averaging the free energies of the different shift models across sessions and participants showed a clear winning model (the instant mapping model), when looking at the winning model separately for sessions within patients, there was some variability. As a post-hoc analysis, the analysis was repeated where we took individual-specific winning models to the group level. Again, no consistent group results were found.

### 3.3 Sensitivity analysis

Even though no consistent group connectivity-pain relations were found, for completeness we present the group results of the sensitivity analysis showing the influence of each connectivity parameter on spectral outcomes in each source. On average, an increase in forward connectivity from the S1-r to the S2-r was related to increases in oscillations between 64 Hz and 71 Hz in the S2-r. In the pIC-r, two clear clusters of increases in oscillations between 17-23 Hz and 28-41 Hz related to changes in connectivity from S2-r to pIC-r were found. In the aIC-r, 6-7 Hz and 10 Hz oscillations were related to connectivity from the pIC-r to the aIC-r. Connectivity from the pIC-l to the aIC-l was related to an increase in oscillations between 13-19 Hz and 22-43 Hz in the aIC-l. Several increases around 28 to 59 Hz oscillations were found in the dACC, which were related to increases in connectivity to these regions from the pIC-l, aIC-r and aIC-l.

Other connections also had an influence on oscillations in the dACC and multiple connections had a relation with oscillations in the frontal regions. Connectivity from the dACC to the DLPFC-r was related to increases between 31-51 Hz in the DLPFC-r. In the mPFC, increased oscillations between 13-15 Hz, 22-23 Hz and 29-45 Hz were found for the connection from the dACC. The full group sensitivity results are displayed in Figure 5.

**Figure 5.**
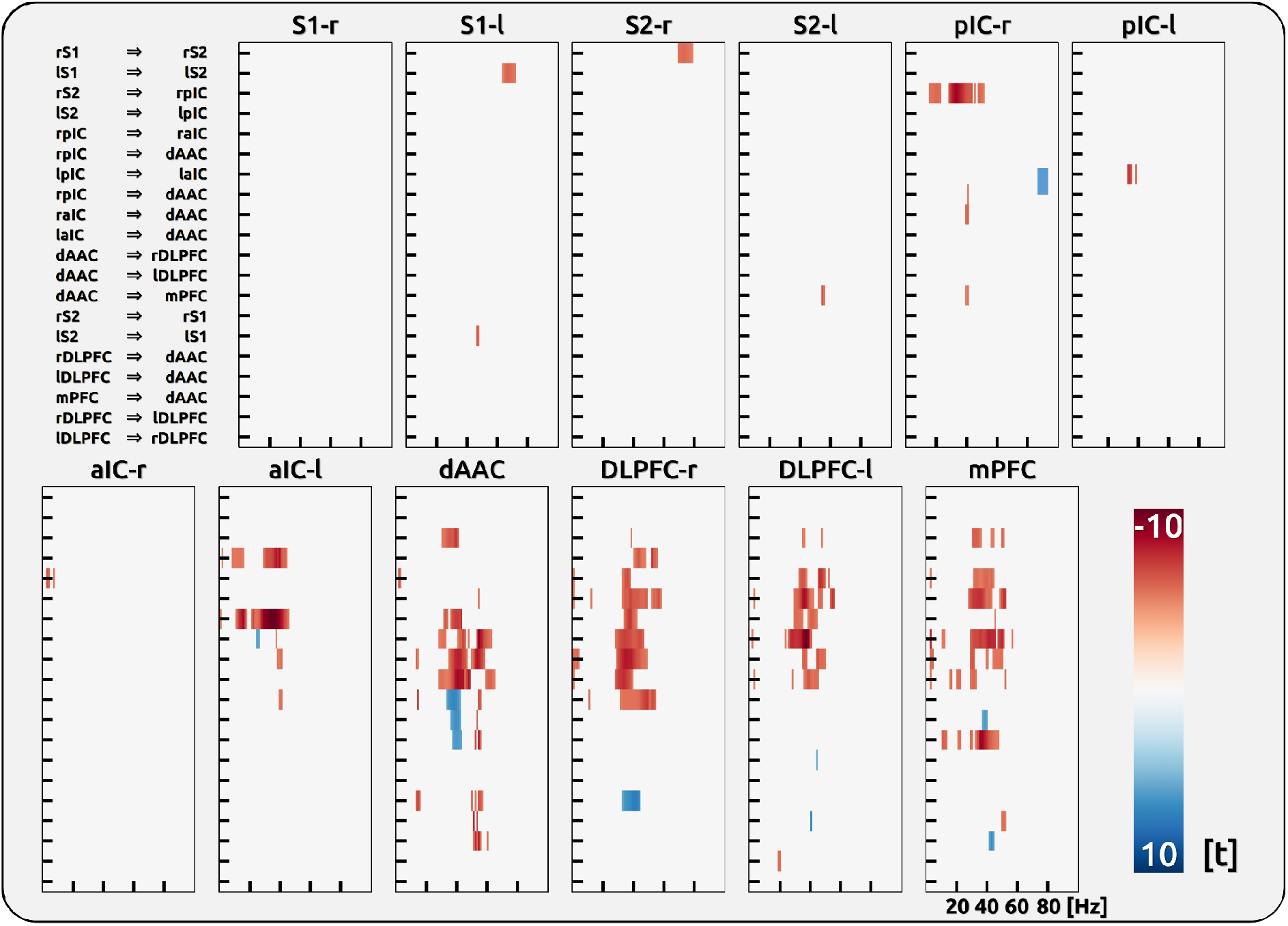
Group results of sensitivity analysis. T values thresholded with the maximum statistic correction. Positive (red) values correspond to connectivity increases resulting in increases in power (at a certain frequency in a specific region), while negative values correspond to decreases in power.

#### 3.3.1 Individual results

The lack of consistency in group results could support the idea of a strong individuality in the relationship between effective connectivity and pain. Besides the group results of the sensitivity analysis that represent some consistent spectral outcomes over patients, connectivity patterns also showed different spectral outcomes depending on the participant. Individual sensitivity profiles can be found in the Supplementary material (Supplementary Figure 3). The average across windows and sessions was taken to compute t-values for visualisation.

## 4 Discussion

Although fluctuating pain is an almost daily experience in CM and other chronic pain conditions, relations between brain dynamics and ongoing pain have been sparsely investigated. Here, we aimed to investigate how dynamic effective connectivity between pain-related regions relates to the fluctuating intensity of ongoing headache in CM. Our pain rating regressors allowed us to examine brain dynamics related to pain intensity (AMP), pain intensity changes (SLP), and pain-unrelated processes such as decision making, motor processing, and changes of visual input (aSLP). At the group level, we found no consistent relations between the connectivity parameters and AMP. In addition, no consistent connectivity changes related to SLP were found. The regressor encoding aSLP showed consistent negative relations with the connections from S1-l to S2-l, S2-l to pIC-l, and aIC-l to dACC, thus mostly encompassing the left ascending pathway. It is not unsurprising that pain-related connections decrease during processes such as decision making, visual and motor processing as connections in other networks might increase, thereby shifting attention from pain (Schulz et al., 2020; Tracey et al., 2002). However, individual data show a more complex picture by suggesting that each patient exhibits their own signature of migraine-related pain encoding in the brain.

### 4.1 Ongoing variations of pain experience in chronic migraine

In a prior publication using the same design and the same patients in fMRI, we applied linear mixed effects models to relate cortical activity and functional connectivity across the whole brain to the same descriptors of the rating process: AMP, SLP and aSLP (Mayr et al., 2021, 2022). Although the data type and analysis strategies differed, the same type of regressors were used to relate brain dynamics to AMP. At group level, several regions and connections were found to be related to AMP and SLP. Importantly, we assessed the similarity between individual cortical results of the CM patients and group results by computing spatial correlations. Variability across individual patterns of connectivity-AMP relations and activity-AMP were large and did not resemble the findings at group level. These previous publications mirror the large individual differences in connectivity-pain relations found in the present study, where we used a linear Bayesian model by taking uncertainties of individual connectivity-pain relations into account.

### 4.2 Preselection of cortical regions

Previous findings of structural and functional abnormalities reported in pain-related areas in patients with migraine (Borsook et al., 2016; Filippi & Messina, 2019; Jia & Yu, 2017; Tolner et al., 2019) supported our motivation to examine the primary and secondary somatosensory cortices, the DLPFC, the medial prefrontal cortex and the dorsal ACC. However, the phenomena investigated in these studies may not be comparable with our findings; most connectivity studies on episodic migraine used a resting state design and tested in the interictal phase during the absence of any headache (Colombo et al., 2015; Maleki & Gollub, 2016; Russo et al., 2017). Although aberrant connectivity between pain-related structures has been found in CM (Hsiao et al., 2021; Lee et al., 2019; Schwedt et al., 2013), the widespread results, methods and study designs makes it difficult to infer hypotheses on the underlying deviant connections. Unfortunately, the demanding computational analysis allows the investigation of only a small number of preselected cortical regions.

### 4.3 Further studies on cortical effects of ongoing variations of pain

There are a few studies that are either investigating applied fluctuating tonic pain in healthy controls or naturally evolving endogenous back pain. For example, a seminal study by Baliki et al. contrasted increasing pain to stable and decreasing chronic back pain which showed higher activity in the right insula, S1, S2, mid cingulate and the cerebellum (Baliki et al., 2006). Further studies found mPFC activity to be related to ongoing pain intensity (Baliki et al., 2006; Hashmi et al., 2013; May et al., 2019; Nickel et al., 2017; Schulz et al., 2015). In particular, gamma oscillations in the mPFC were found to be related to subjective pain intensity of both tonic pain (Nickel et al., 2017; Schulz et al., 2015) and chronic back pain (May et al., 2019). Connectivity at alpha frequencies in the sensorimotor-prefrontal network has also been shown to be associated with tonic pain in healthy controls (Nickel et al., 2020). Based on these studies, we would have expected some results containing connectivity parameters from and to the mPFC. However, these previous studies were analysing the amplitude of cortical processes in different samples; they are not directly comparable to the present variable connectivity scores in migraine patients. Furthermore, mPFC activity being related to subjective tonic or chronic back pain intensity may not be consistent across subjects. In the supplementary material of the study of Schulz et al. (Schulz et al., 2015), some participants indeed showed clear positive relations between prefrontal gamma oscillations and pain intensity but others showed no or even negative relations. In a similar study on chronic back pain (May et al., 2019), group data showed a positive relation between ongoing pain and gamma oscillations at the Fz electrode, but some patients showed no or negative relations. From the 31 patients, only 9 showed a positive relation that had a p-value lower than 0.1.

### 4.4 Individual signatures of AMP encoding in migraine using DCM

Therefore, the most important question we have to address is whether this study was able to reveal brain dynamics related to pain in individual patients. Each patient underwent four recording sessions to investigate consistency of pain processing patterns. Across all sessions, 16 of the 20 patients showed one or more consistent relation with some connectivity parameter in the pain network and AMP and 17 patients showed one or more consistent relation with falling and rising pain. The variable results do not just encompass differences in the magnitude of relations between connectivity and pain processing across individuals, but also in the directionality of the relations. For example some patients show enhanced connectivity from region A to B when their pain increases, whereas others show decreased connectivity. The interpretation of these findings depends mostly on the function(s) encoded by that region and whether that region can have both inhibitory and excitatory connections with other regions. Variability in the sensitivity profiles do suggest that a change in connectivity between two regions may have different spectral outcomes depending on the patient. Mayr et al. (Mayr et al., 2022) also found qualitative differences across patients in relations between brain connectivity and AMP. These findings open the question whether there are mechanisms that determine a positive, negative, or non-existent relation between brain activity and AMP for a given connection.

### 4.5 Requirements for the applicability of individual descriptors brain dynamics

The current study design was aimed to approximate the daily-life experience of migraine patients, which is the experience of fluctuating pain intensity. The assessment of repeated sessions is mandatory in order to minimise the influence of random noise that may occur within single sessions. To better understand the inherently dynamic individual pain experience, it is important to further investigate the stability of the relevant pain-related cortical processes over time (Mun et al., 2019). Similar to our previous study (Mayr et al., 2022), we suggest that individual-specific neural reorganisation due to repeated attacks could underlie qualitative rather than gradual/quantitative differences between individuals. Such reliable cortical target processes could be utilised to accurately predict the pain ratings in each patient. In a follow-up study, we will use machine learning techniques to explore whether pain ratings can be predicted on the basis of the individual brain dynamics that we have analysed in the present investigation. We would assume a better prediction at the individual level compared to predictions that are based on data from the entire sample. In a similar vein, we expect a stronger future focus on individual analyses (Martucci et al., 2014). Consequently, determining individually unique and stable brain dynamics that encode AMP and SLP in single patients could be a prerequisite to define a reliable target for possible neuromodulatory interventions (Jensen et al., 2014).

### 4.6 Limitations

This study was a first step to gain insight in relations between fluctuating connectivity and the almost daily headaches that CM patients experience. Thus, we began with testing the cortical networks most mentioned in the pain literature. Nevertheless, there is a possibility that we have not selected the network most important for pain processing in CM.

In addition, other models might be better suited to investigate the dynamically evolving pain-related oscillations. Sophisticated methods like hidden Markov models (Hughey & Krogh, 1996) could be used to examine transitions of different brain states and how these states relate to the reported pain. Auto-regressive models on the pain ratings and comparing models with different regressors of smoothed fitted ratings could gain further insight.

Finally, the present design does not allow to include a healthy control group. This group would likely show some fluctuating network activity. However, the pain rating would consist of a rating time course of zeros. A “correlation” with zeros is mathematically not possible and a healthy control group using this design is therefore not possible.

### 4.7 Summary and outlook

The current findings support other studies from our research group showing that individual patients have unique patterns of brain dynamics underlying their chronic pain that are difficult to capture. Using neuroimaging to find a biomarker for subjective pain intensity at the group level might not lead to fruitful results. Finding the appropriate neuroimaging method and analysis to establish brain markers that are stable throughout multiple recording sessions of the same patient may be the way forward to guide individualised treatments. Finding subtypes of patients that process pain in the same way would be valuable but would require a large sample of patients, each with repeated recordings.

## Supporting information

All patients were permitted to continue their pharmacological treatment at a stable dose

Individual sensitivity profiles can be found in the Supplementary material

## Acknowledgements

This work was supported by the Fund for Scientific Research-Flanders (FWO-V, PhD Fellowship [grant number FWO17/ASP/042] awarded to IB), the Special Research Funds of Ghent University (DM and FVDS), and by the German Research Fund awarded to ES (DFG, SCHU 2879/4-1). We thank Dr Stephanie Irving for copy-editing the manuscript.

