## Supplementary material for "Investigation on How Dynamic Effective Connectivity Patterns Encode the Fluctuating Pain Intensity in Chronic Migraine": All patients were permitted to continue their pharmacological treatment at a stable dose

### Supplement

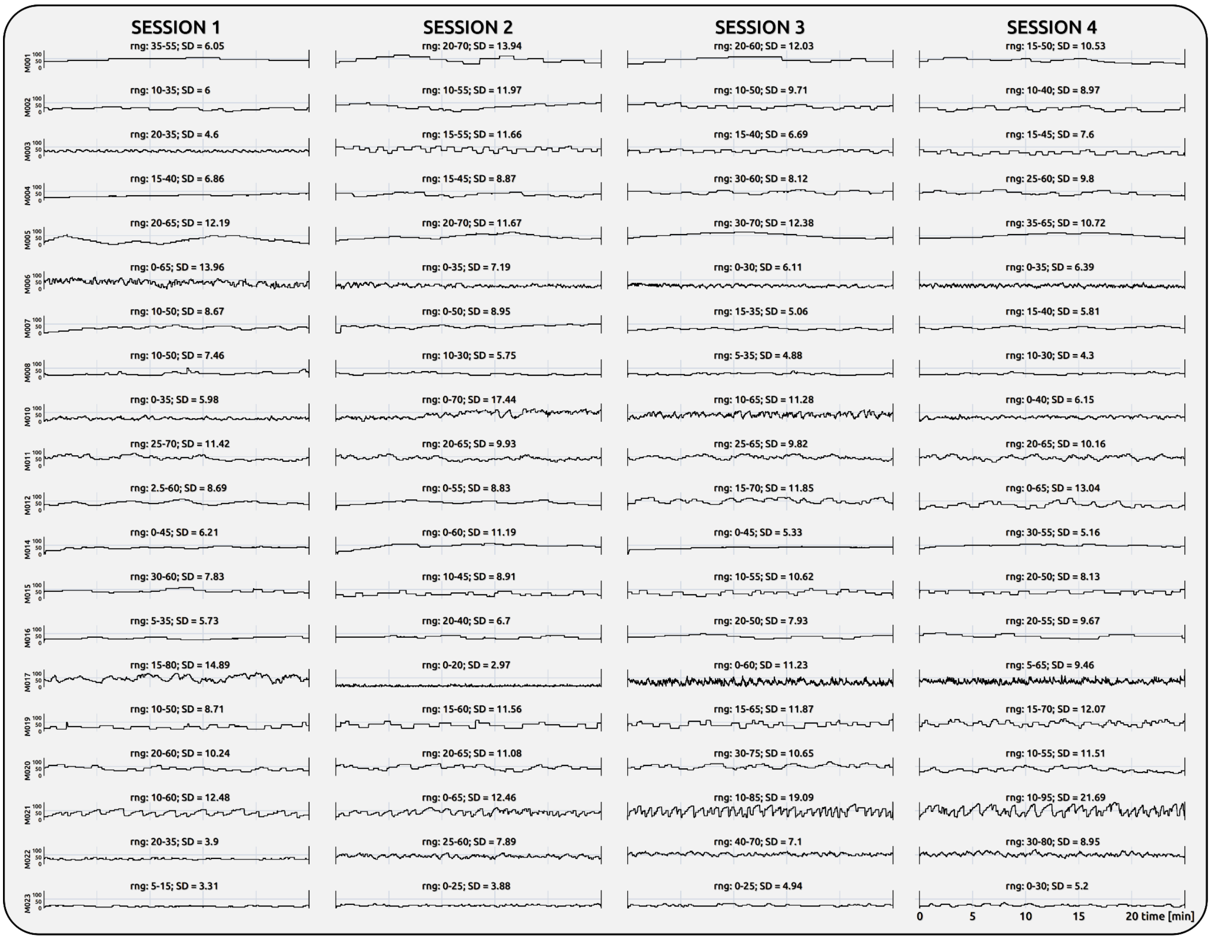


**Supplementary Figure 1 |** The figure shows the time course of pain intensity (AMP) across the 25 min experiment. The range (rng) of the pain intensity rating (min-max) and the standard deviation (SD) over the whole time series are specified.

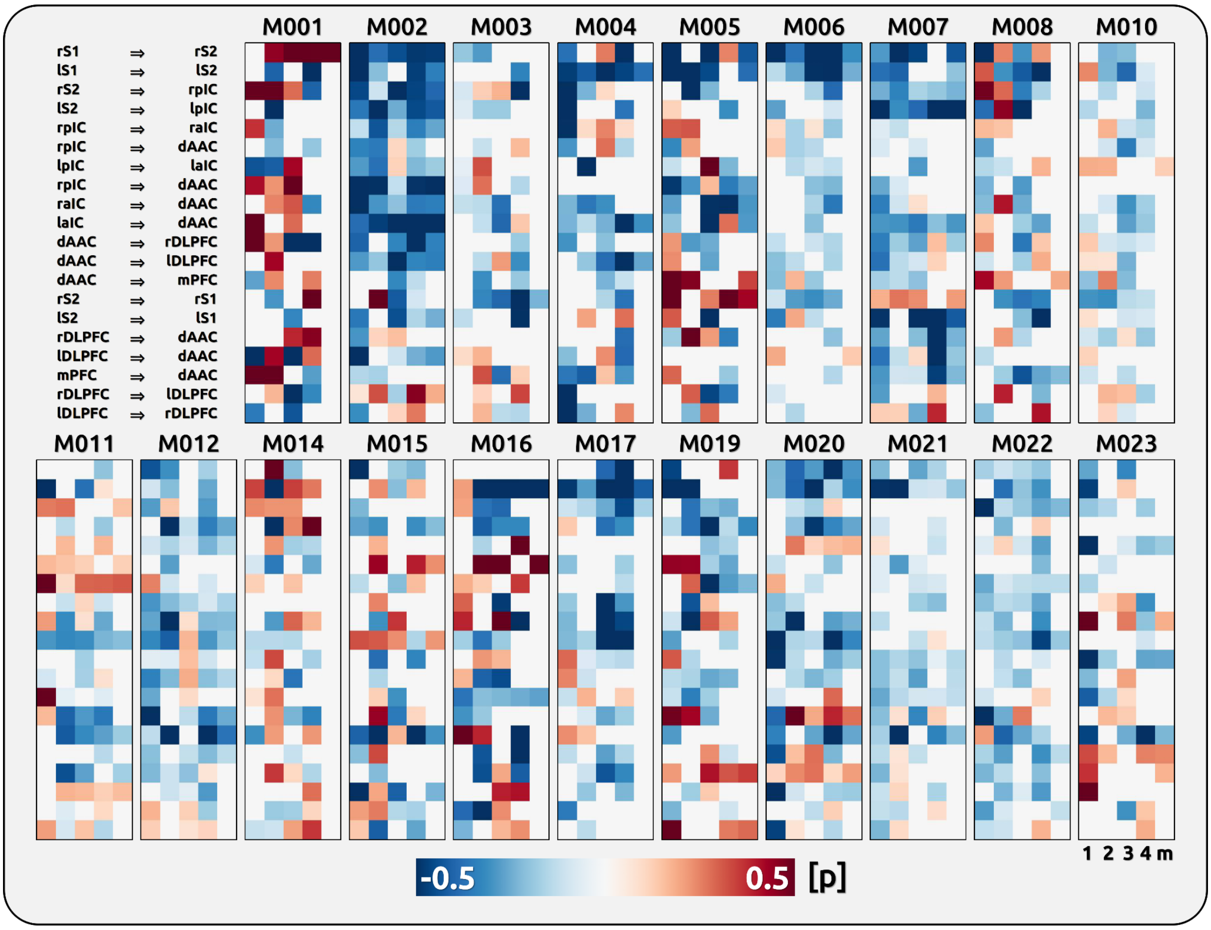


**Supplementary Figure 2 | Individual results of relations between the connectivity parameters and aSLP in each session and across sessions.** Columns 1 to 4 represent the beta values from the within-session level PEB and are thresholded at a posterior probability > 0.75. Column 5 represents the beta values from the within-subject PEB, thresholded at a posterior probability > 0.95.

**Supplementary Table 1 | Characteristics of CM patients reported in questionnaires**

| # | m/f | age (years) | pain duration (years) | pain medication | pain intensity | PCS | d/a/s |
|---|---|---|---|---|---|---|---|
| 1 | f | 61 | 50 | Sumatriptan 100mg (20x/month) | 7 | 15 | 0/10/9 |
| 2 | f | 27 | 7 | Metamizole 500mg (2-3x/month), Sumatriptan 50mg (1x/month) | 4 | 5 | 0/0/2 |
| 3 | f | 50 | 35 | Sumatriptan 100mg (5-7x/month) | 4 | 24 | 5/0/6 |
| 4 | f | 27 | 8 | Zolmitriptan 20mg (2x/month), Ibuprofen 600mg (7x/month) | 7 | 37 | 1/1/8 |
| 5 | m | 49 | 30 | Ibuprofen 600mg (7-8x/month), Metamizole 500mg (3-4x/month), Paracetamol 500mg (5-6x/month) | 4 | 3 | 0/6/1 |
| 6 | f | 52 | 30 | Ibuprofen 400mg (6x/month), Paracetamol 1000mg (2x/month) | 5 | 11 | 5/10/12 |
| 7 | f | 32 | 15 | Zolmitriptan 5mg (8x/month), Naproxen 500mg (15x/month), Acetylsalicylic Acid (ASA) 250mg (4x/month), Paracetamol 200mg (4x/month), Caffeine 50mg (4x/month) | 4 | 31 | 5/7/7 |
| 8 | f | 21 | 7 | Sumatriptan 50mg (1x/month) | 4 | 10 | 2/1/0 |
| 9 | f | 19 | 7 | none | 4 | 35 | 10/2/6 |
| 10 | f | 46 | 13 | Ibuprofen 800mg (8-10x/month) | 6 | 31 | 7/10/15 |
| 11 | f | 27 | 13 | Triptan (2-3x/month), ASA 250mg (20-25x/month), Paracetamol 250mg (20-25x/month), Caffeine 50mg (20-25x/month) | 4 | 13 | 6/1/10 |
| 12 | m | 53 | 15 | Ibuprofen 600mg (10x/month) | 6 | 20 | 7/2/4 |
| 13 | f | 30 | 6 | Ibuprofen 400mg (4x/month) | 4 | 24 | 1/1/5 |
| 14 | f | 21 | 7 | Ibuprofen 600mg (4-8x/month), Paracetamol 500mg (4x/month), Zolmitriptan 5mg (1-2x/month) | 3 | 15 | 2/0/5 |
| 15 | f | 23 | 8 | Ibuprofen 600mg (2-3x/month) | 7 | 24 | 2/0/3 |
| 16 | f | 28 | 7 | Ibuprofen 500mg (10x/month) | 7 | 11 | 0/1/1 |
| 17 | f | 25 | 5 | Ibuprofen 600mg (5x/month), Zolmitriptan 5mg (1x/month) | 4 | 21 | 5/5/9 |
| 18 | f | 33 | 20 | Paracetamol 500mg (3x/month) | 5 | 32 | 4/0/4 |
| 19 | f | 21 | 9 | Ibuprofen 400mg (6-10x/month), Rizatriptan 10mg (2x/month) | 5 | 30 | 1/1/2 |
| 20 | f | 43 | 10 | Ibuprofen 400mg (20x/month), Paracetamol 325 mg (8-10x/month), Naproxen 100 mg (8-10x/month), Caffeine 50 mg (8-10x/month), Drotaverine hydrochloride 40 mg (8-10x/month), Pheniramine 10 mg (8-10x/month) | 4 | 22 | 2/3/9 |

m/f: male/female; PCS: pain catastrophizing scale; d/a/s: depression/anxiety/stress. The cutoff for depression and stress is 10, for anxiety 6, and for the PCS 30.

**Supplementary Table 2 |. MNI coordinates of the included brain regions**

| brain region | x | y | z |
|---|---|---|---|
| right primary somatosensory cortex (rS1) | 52 | -20 | 38 |
| left primary somatosensory cortex (lS1) | -58 | -20 | 36 |
| right secondary somatosensory cortex (rS2) | 56 | -24 | 22 |
| left secondary somatosensory cortex (lS2) | -58 | -23 | 19 |
| right anterior insula (raIC) | 38 | 8 | 4 |
| left anterior insula (laIC) | -40 | 8 | -6 |
| right posterior insula (rpIC) | 38 | -16 | 12 |
| left posterior insula (lpIC) | -36 | -18 | 11 |
| right dorsolateral prefrontal cortex (rDLPFC) | 42 | 35 | 26 |
| left dorsolateral prefrontal cortex (lDLPFC) | -39 | 29 | 26 |
| dorsal anterior cingulate cortex (dACC) | 0 | 8 | 36 |
