## Supplementary material for "Investigation on How Dynamic Effective Connectivity Patterns Encode the Fluctuating Pain Intensity in Chronic Migraine": Individual sensitivity profiles can be found in the Supplementary material

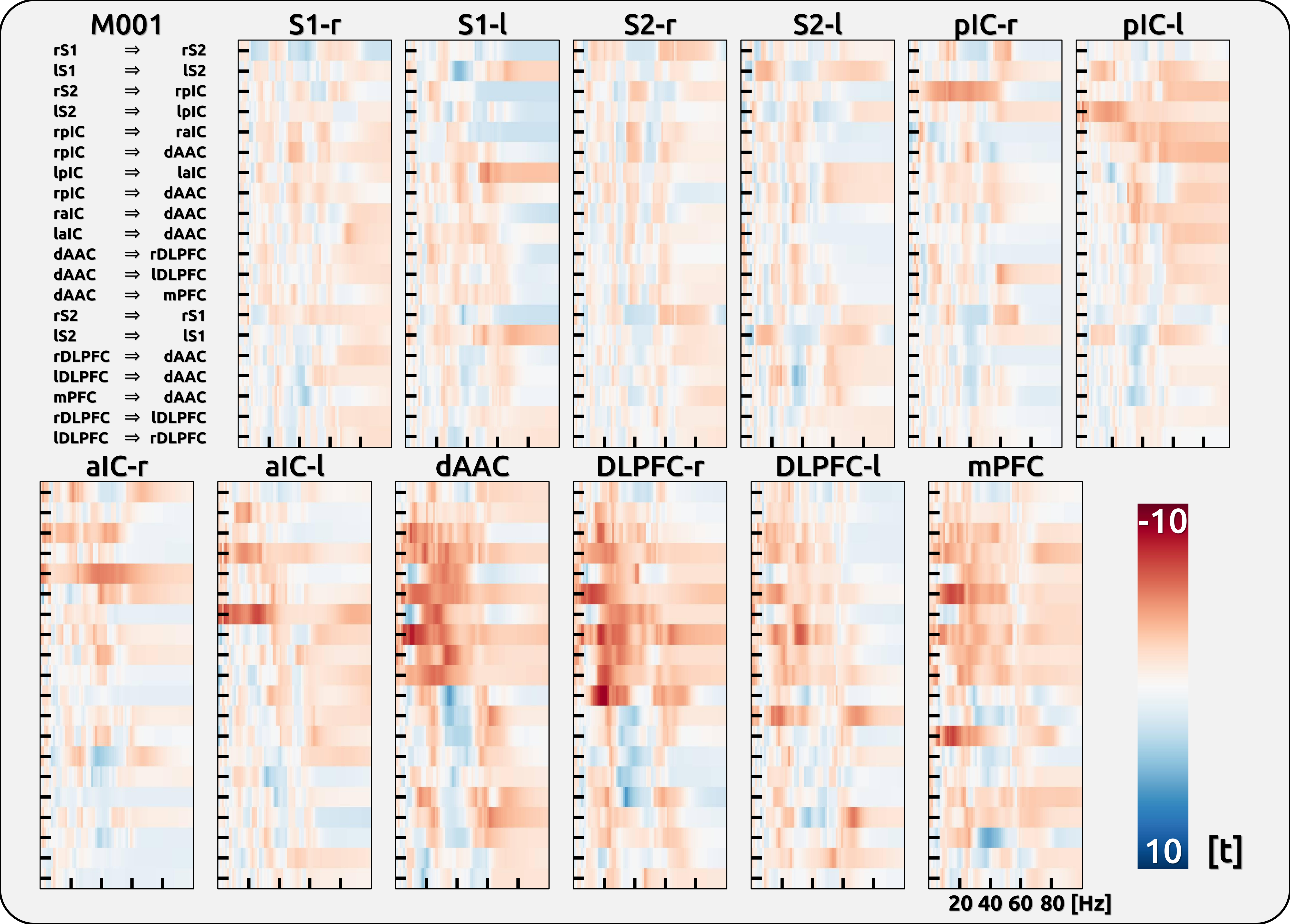
M001
rS1 ⇒ rS2
lS1 ⇒ lS2
rS2 ⇒ rpIC
lS2 ⇒ lpIC
rpIC ⇒ raIC
rpIC ⇒ dAAC
lpIC ⇒ laIC
rpIC ⇒ dAAC
raIC ⇒ dAAC
laIC ⇒ dAAC
dAAC ⇒ rDLPFC
dAAC ⇒ lDLPFC
dAAC ⇒ mPFC
rS2 ⇒ rS1
lS2 ⇒ lS1
rDLPFC ⇒ dAAC
lDLPFC ⇒ dAAC
mPFC ⇒ dAAC
rDLPFC ⇒ lDLPFC
lDLPFC ⇒ rDLPFC
S1-r
S1-l
S2-r
S2-l
pIC-r
pIC-l
aIC-r
aIC-l
dAAC
DLPFC-r
DLPFC-l
mPFC
-10
10
[t]
20 40 60 80 [Hz]

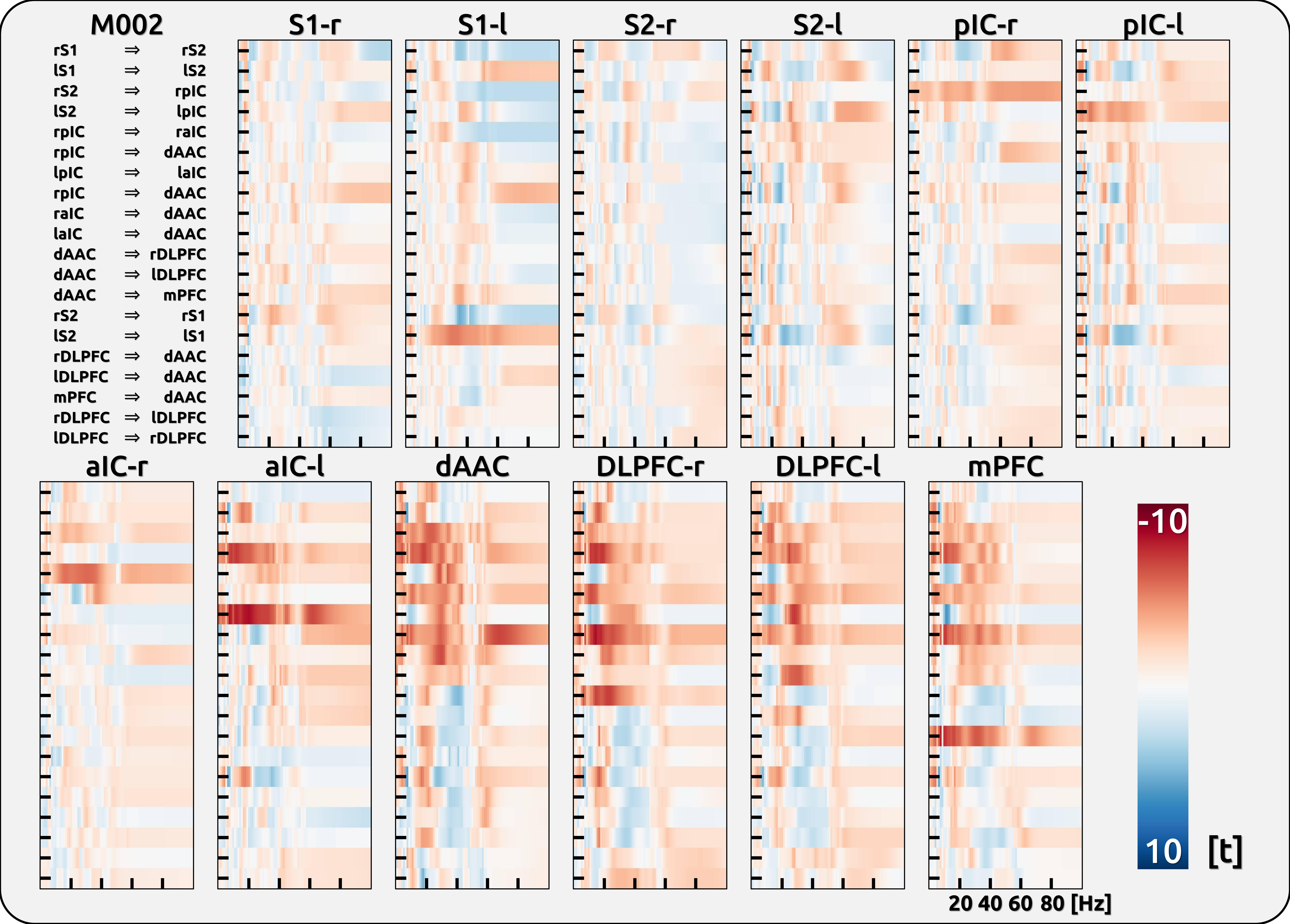

M002
S1-r
S1-l
S2-r
S2-l
pIC-r
pIC-l
rS1 ⇒ rS2
lS1 ⇒ lS2
rS2 ⇒ rpIC
lS2 ⇒ lpIC
rpIC ⇒ raIC
rpIC ⇒ dAAC
lpIC ⇒ laIC
rpIC ⇒ dAAC
raIC ⇒ dAAC
laIC ⇒ dAAC
dAAC ⇒ rDLPFC
dAAC ⇒ lDLPFC
dAAC ⇒ mPFC
rS2 ⇒ rS1
lS2 ⇒ lS1
rDLPFC ⇒ dAAC
lDLPFC ⇒ dAAC
mPFC ⇒ dAAC
rDLPFC ⇒ lDLPFC
lDLPFC ⇒ rDLPFC
aIC-r
aIC-l
dAAC
DLPFC-r
DLPFC-l
mPFC
-10
10
[t]
20 40 60 80 [Hz]

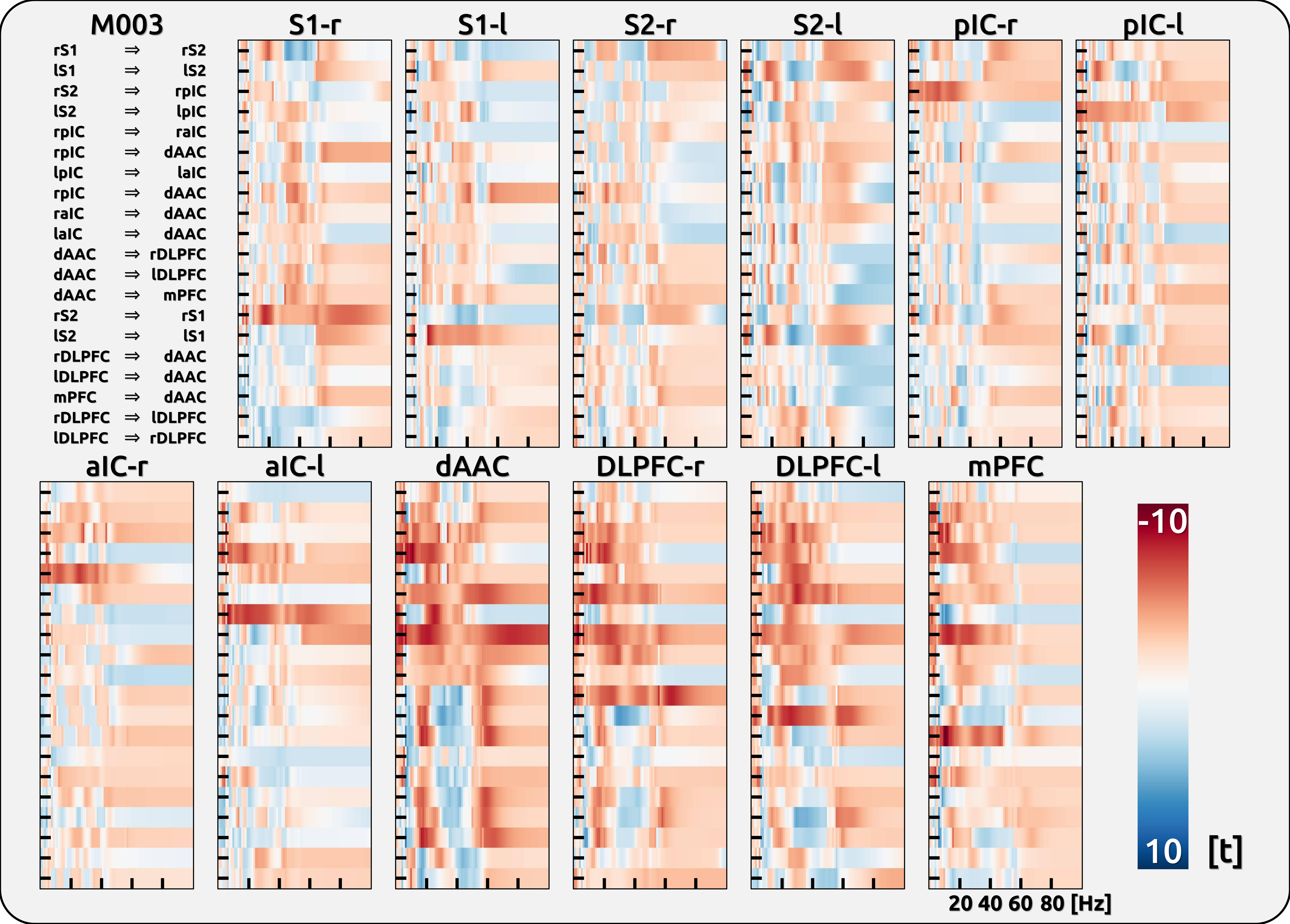
M003
rS1 ⇒ rS2
lS1 ⇒ lS2
rS2 ⇒ rpIC
lS2 ⇒ lpIC
rpIC ⇒ raIC
rpIC ⇒ dAAC
lpIC ⇒ laIC
rpIC ⇒ dAAC
raIC ⇒ dAAC
laIC ⇒ dAAC
dAAC ⇒ rDLPFC
dAAC ⇒ lDLPFC
dAAC ⇒ mPFC
rS2 ⇒ rS1
lS2 ⇒ lS1
rDLPFC ⇒ dAAC
lDLPFC ⇒ dAAC
mPFC ⇒ dAAC
rDLPFC ⇒ lDLPFC
lDLPFC ⇒ rDLPFC
S1-r
S1-l
S2-r
S2-l
pIC-r
pIC-l
aIC-r
aIC-l
dAAC
DLPFC-r
DLPFC-l
mPFC
-10
10
[t]
20 40 60 80 [Hz]

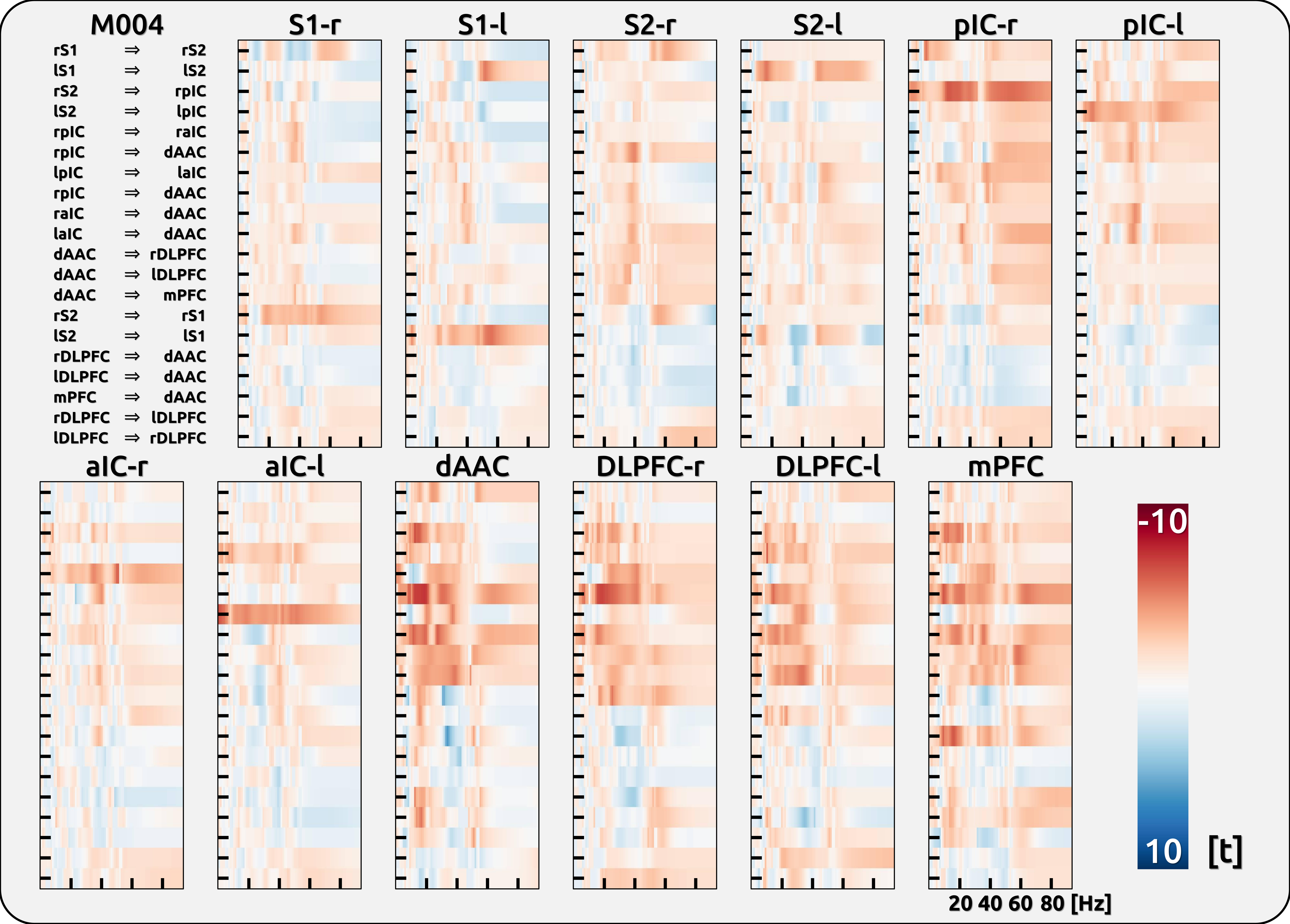

M004
S1-r
S1-l
S2-r
S2-l
pIC-r
pIC-l
rS1 ⇒ rS2
lS1 ⇒ lS2
rS2 ⇒ rpIC
lS2 ⇒ lpIC
rpIC ⇒ raIC
rpIC ⇒ dAAC
lpIC ⇒ laIC
rpIC ⇒ dAAC
raIC ⇒ dAAC
laIC ⇒ dAAC
dAAC ⇒ rDLPFC
dAAC ⇒ lDLPFC
dAAC ⇒ mPFC
rS2 ⇒ rS1
lS2 ⇒ lS1
rDLPFC ⇒ dAAC
lDLPFC ⇒ dAAC
mPFC ⇒ dAAC
rDLPFC ⇒ lDLPFC
lDLPFC ⇒ rDLPFC
aIC-r
aIC-l
dAAC
DLPFC-r
DLPFC-l
mPFC
-10
10
[t]
20 40 60 80 [Hz]

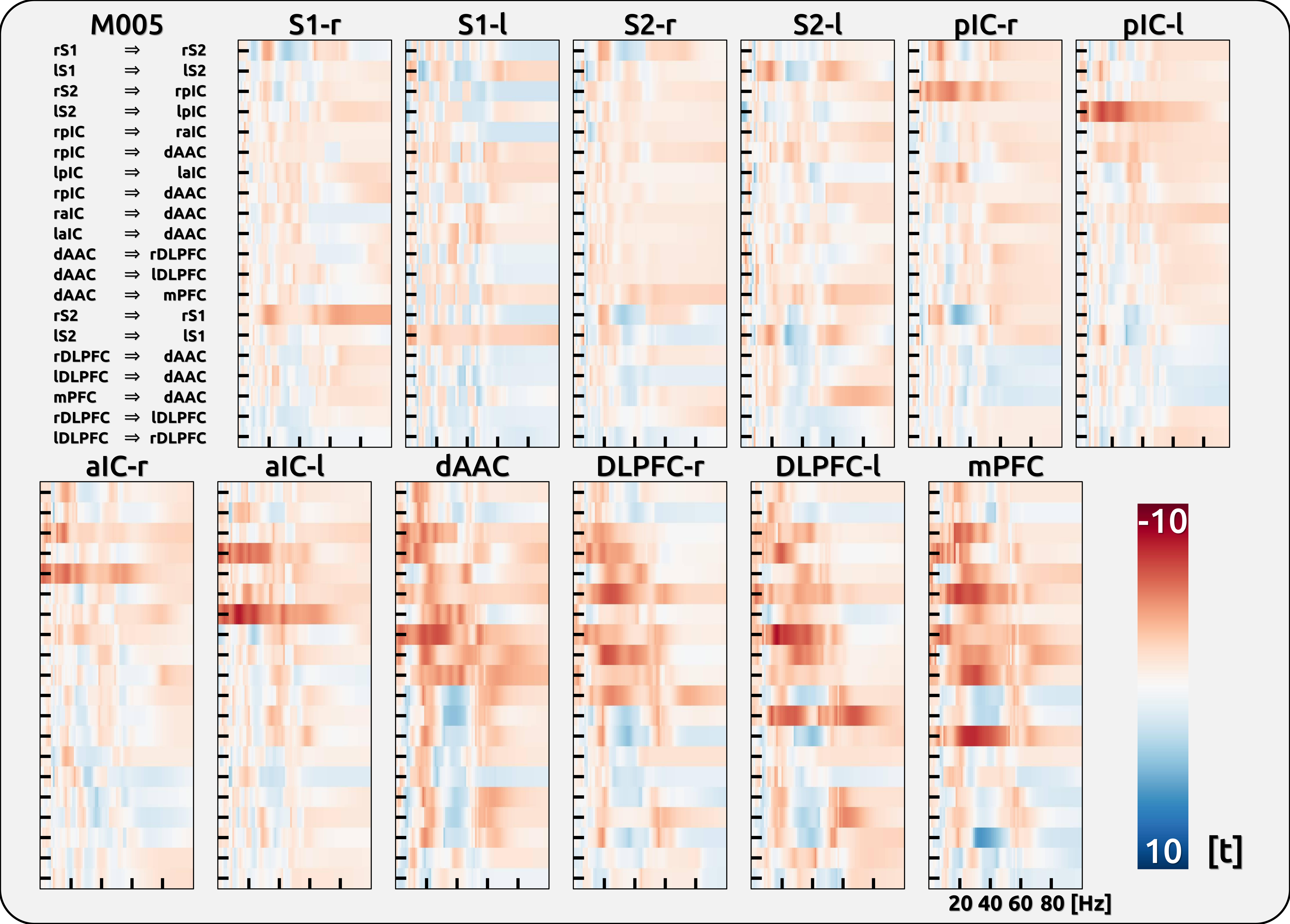

M005
rS1 ⇒ rS2
lS1 ⇒ lS2
rS2 ⇒ rpIC
lS2 ⇒ lpIC
rpIC ⇒ raIC
rpIC ⇒ dAAC
lpIC ⇒ laIC
rpIC ⇒ dAAC
raIC ⇒ dAAC
laIC ⇒ dAAC
dAAC ⇒ rDLPFC
dAAC ⇒ lDLPFC
dAAC ⇒ mPFC
rS2 ⇒ rS1
lS2 ⇒ lS1
rDLPFC ⇒ dAAC
lDLPFC ⇒ dAAC
mPFC ⇒ dAAC
rDLPFC ⇒ lDLPFC
lDLPFC ⇒ rDLPFC
S1-r
S1-l
S2-r
S2-l
pIC-r
pIC-l
aIC-r
aIC-l
dAAC
DLPFC-r
DLPFC-l
mPFC
-10
10
[t]
20 40 60 80 [Hz]

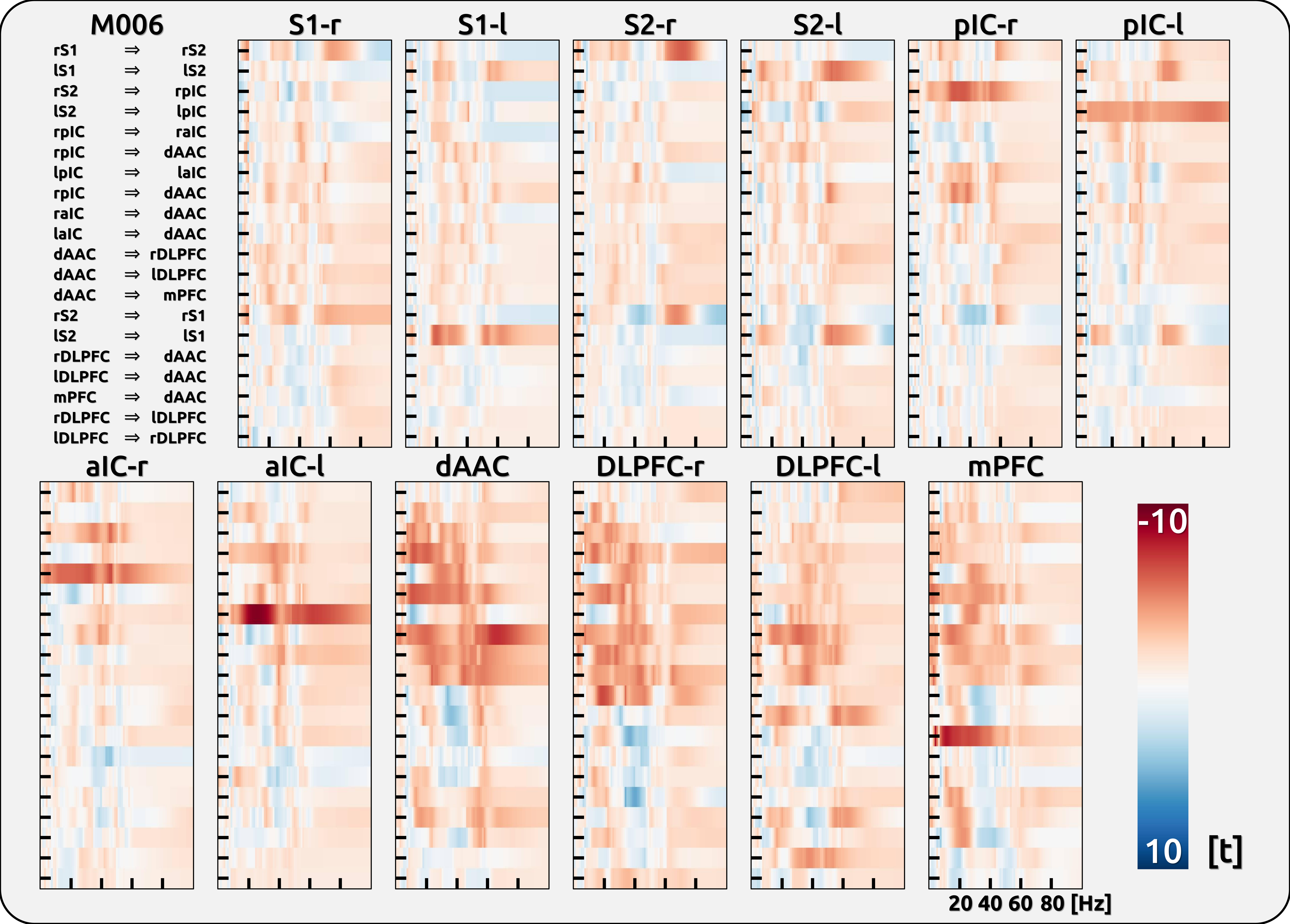
M006
S1-r
S1-l
S2-r
S2-l
pIC-r
pIC-l
rS1 ⇒ rS2
lS1 ⇒ lS2
rS2 ⇒ rpIC
lS2 ⇒ lpIC
rpIC ⇒ raIC
rpIC ⇒ dAAC
lpIC ⇒ laIC
rpIC ⇒ dAAC
raIC ⇒ dAAC
laIC ⇒ dAAC
dAAC ⇒ rDLPFC
dAAC ⇒ lDLPFC
dAAC ⇒ mPFC
rS2 ⇒ rS1
lS2 ⇒ lS1
rDLPFC ⇒ dAAC
lDLPFC ⇒ dAAC
mPFC ⇒ dAAC
rDLPFC ⇒ lDLPFC
lDLPFC ⇒ rDLPFC
aIC-r
aIC-l
dAAC
DLPFC-r
DLPFC-l
mPFC
-10
10
[t]
20 40 60 80 [Hz]

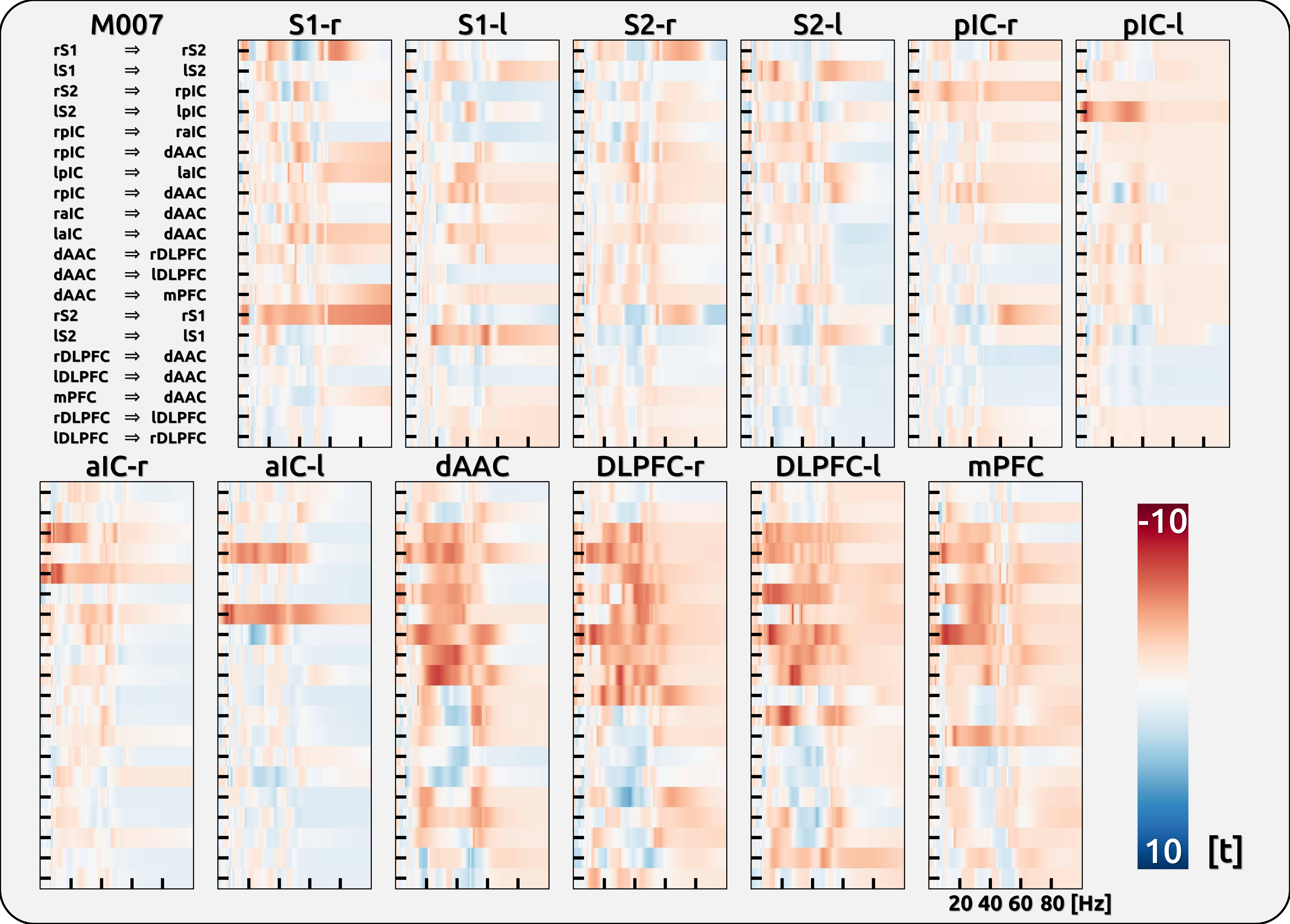
M007
S1-r
S1-l
S2-r
S2-l
pIC-r
pIC-l
rS1 ⇒ rS2
lS1 ⇒ lS2
rS2 ⇒ rpIC
lS2 ⇒ lpIC
rpIC ⇒ raIC
rpIC ⇒ dAAC
lpIC ⇒ laIC
rpIC ⇒ dAAC
raIC ⇒ dAAC
laIC ⇒ dAAC
dAAC ⇒ rDLPFC
dAAC ⇒ lDLPFC
dAAC ⇒ mPFC
rS2 ⇒ rS1
lS2 ⇒ lS1
rDLPFC ⇒ dAAC
lDLPFC ⇒ dAAC
mPFC ⇒ dAAC
rDLPFC ⇒ lDLPFC
lDLPFC ⇒ rDLPFC
aIC-r
aIC-l
dAAC
DLPFC-r
DLPFC-l
mPFC
-10
10
[t]
20 40 60 80 [Hz]

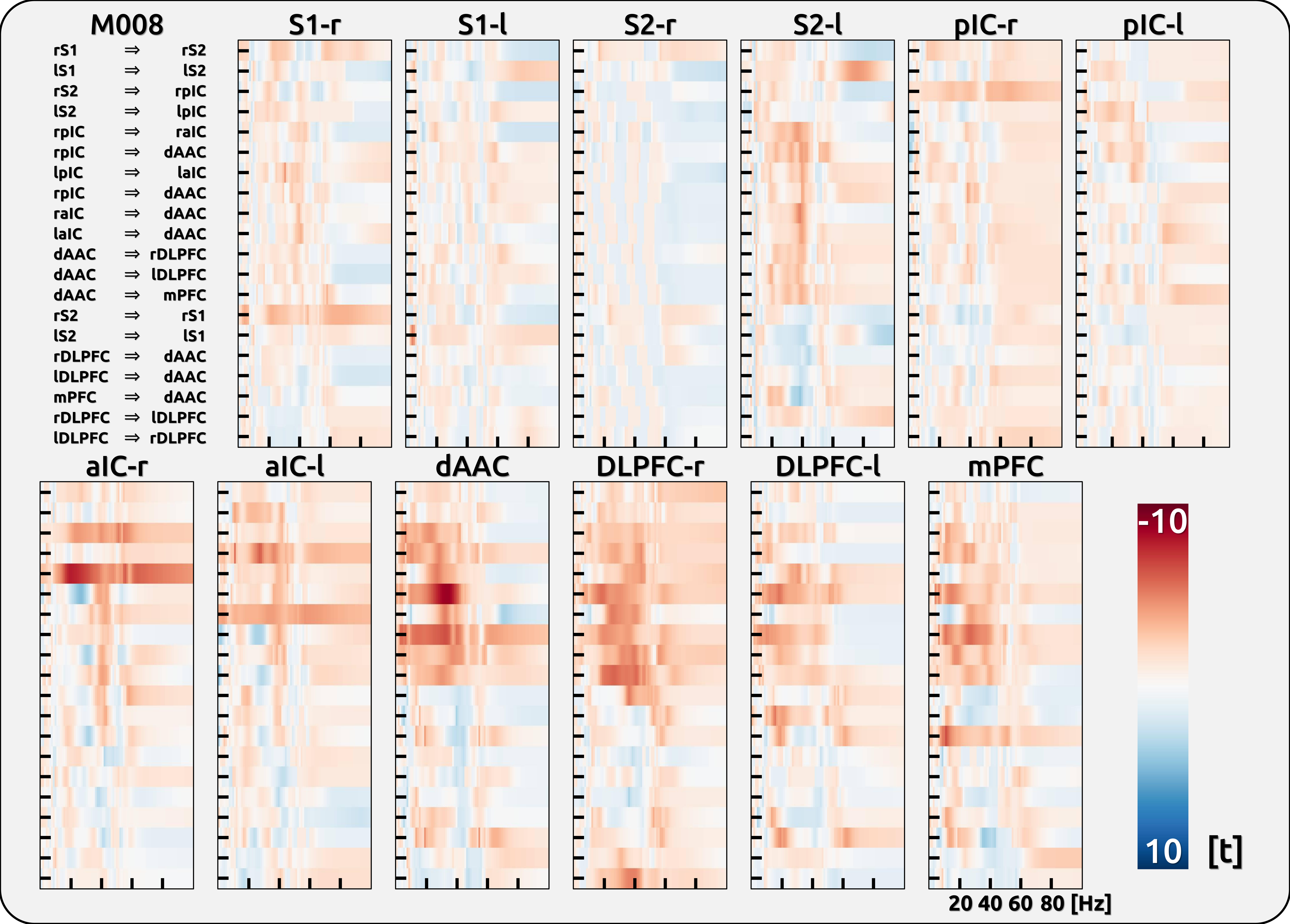

M008
S1-r
S1-l
S2-r
S2-l
pIC-r
pIC-l
rS1 ⇒ rS2
lS1 ⇒ lS2
rS2 ⇒ rpIC
lS2 ⇒ lpIC
rpIC ⇒ raIC
rpIC ⇒ dAAC
lpIC ⇒ laIC
rpIC ⇒ dAAC
raIC ⇒ dAAC
laIC ⇒ dAAC
dAAC ⇒ rDLPFC
dAAC ⇒ lDLPFC
dAAC ⇒ mPFC
rS2 ⇒ rS1
lS2 ⇒ lS1
rDLPFC ⇒ dAAC
lDLPFC ⇒ dAAC
mPFC ⇒ dAAC
rDLPFC ⇒ lDLPFC
lDLPFC ⇒ rDLPFC
aIC-r
aIC-l
dAAC
DLPFC-r
DLPFC-l
mPFC
-10
10
[t]
20 40 60 80 [Hz]

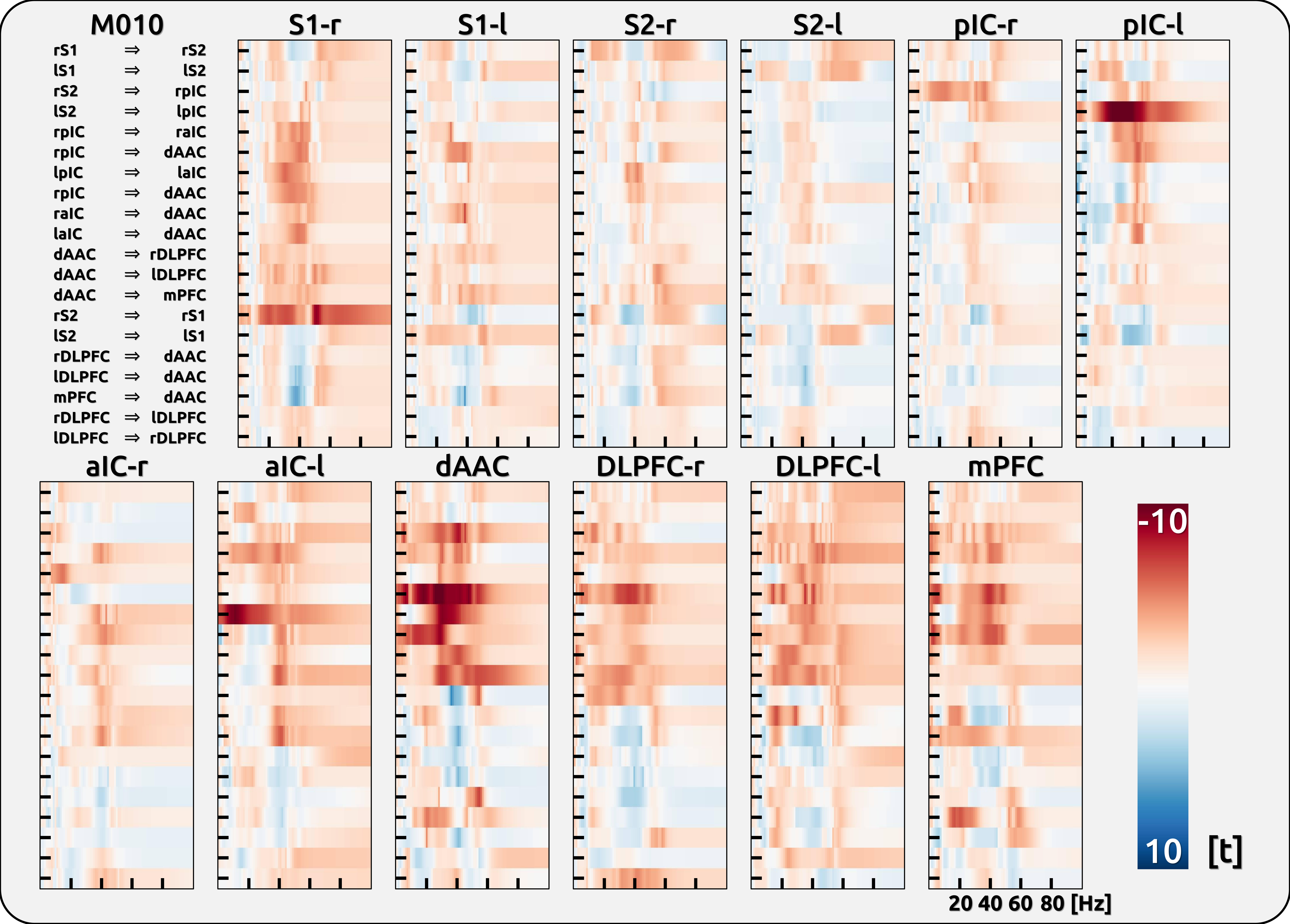

M010
rS1 ⇒ rS2
lS1 ⇒ lS2
rS2 ⇒ rpIC
lS2 ⇒ lpIC
rpIC ⇒ raIC
rpIC ⇒ dAAC
lpIC ⇒ laIC
rpIC ⇒ dAAC
raIC ⇒ dAAC
laIC ⇒ dAAC
dAAC ⇒ rDLPFC
dAAC ⇒ lDLPFC
dAAC ⇒ mPFC
rS2 ⇒ rS1
lS2 ⇒ lS1
rDLPFC ⇒ dAAC
lDLPFC ⇒ dAAC
mPFC ⇒ dAAC
rDLPFC ⇒ lDLPFC
lDLPFC ⇒ rDLPFC
S1-r
S1-l
S2-r
S2-l
pIC-r
pIC-l
aIC-r
aIC-l
dAAC
DLPFC-r
DLPFC-l
mPFC
-10
10
[t]
20 40 60 80 [Hz]

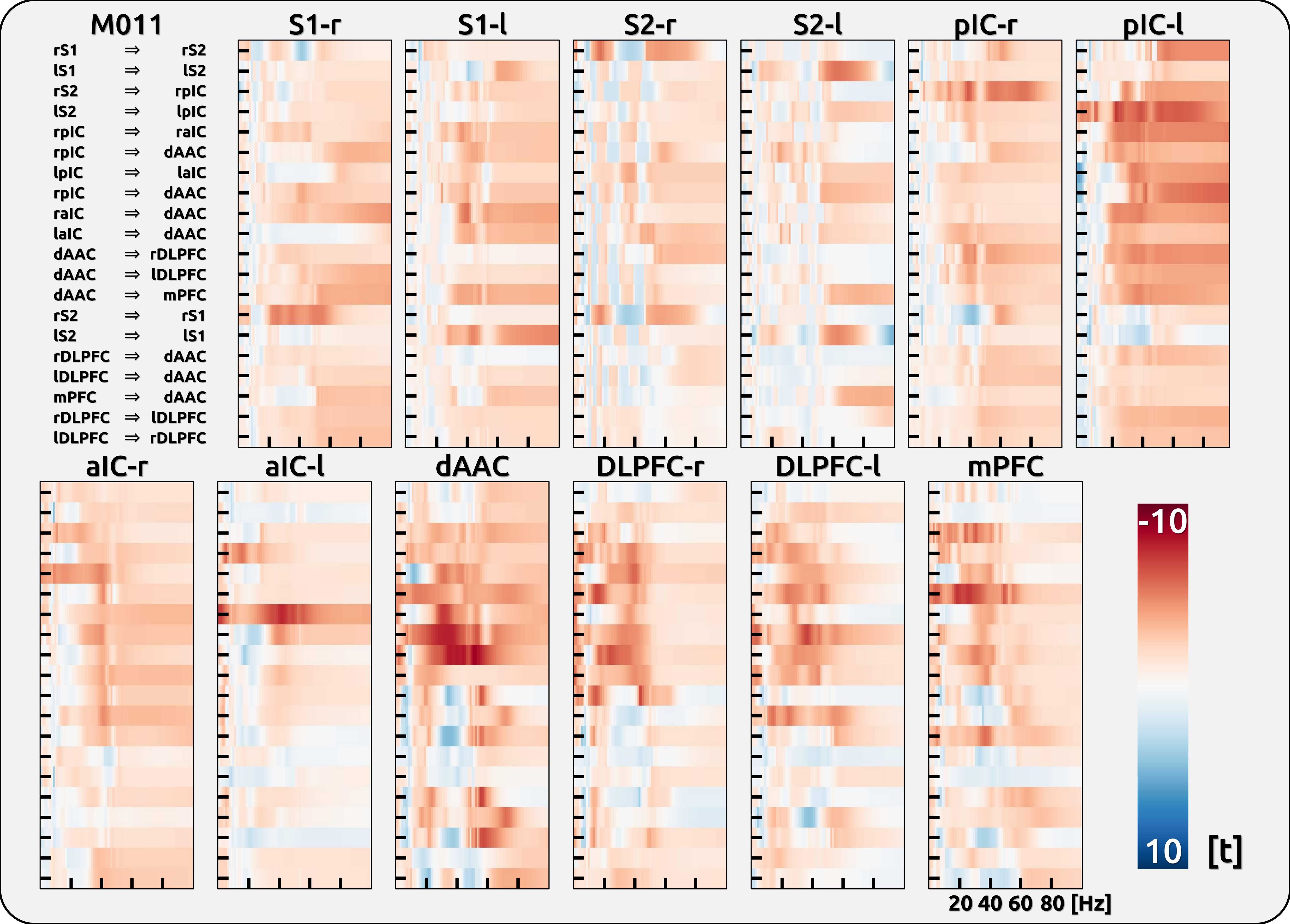

M011
rS1 ⇒ rS2
lS1 ⇒ lS2
rS2 ⇒ rpIC
lS2 ⇒ lpIC
rpIC ⇒ raIC
rpIC ⇒ dAAC
lpIC ⇒ laIC
rpIC ⇒ dAAC
raIC ⇒ dAAC
laIC ⇒ dAAC
dAAC ⇒ rDLPFC
dAAC ⇒ lDLPFC
dAAC ⇒ mPFC
rS2 ⇒ rS1
lS2 ⇒ lS1
rDLPFC ⇒ dAAC
lDLPFC ⇒ dAAC
mPFC ⇒ dAAC
rDLPFC ⇒ lDLPFC
lDLPFC ⇒ rDLPFC
S1-r
S1-l
S2-r
S2-l
pIC-r
pIC-l
aIC-r
aIC-l
dAAC
DLPFC-r
DLPFC-l
mPFC
-10
10
[t]
20 40 60 80 [Hz]

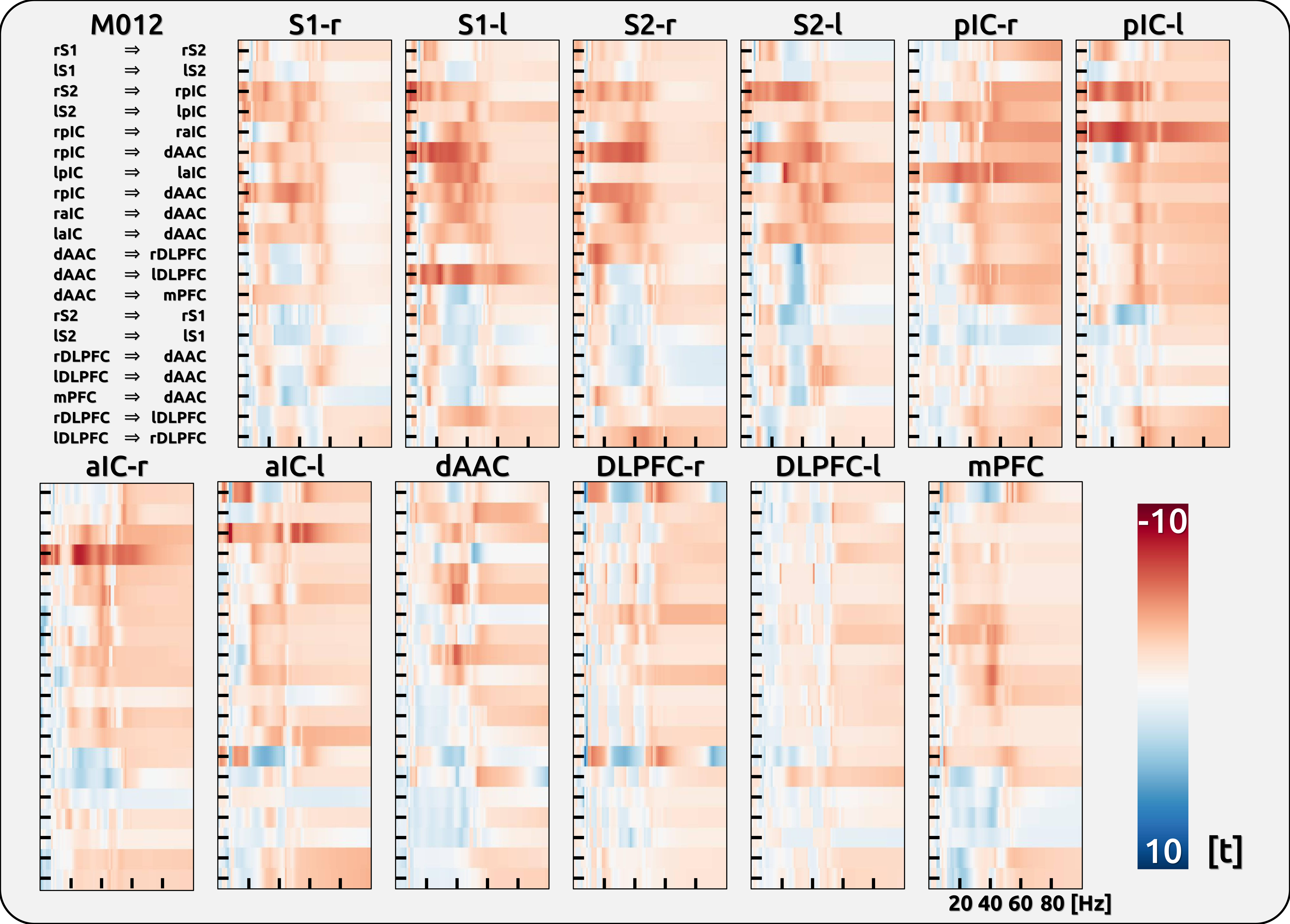

M012
rS1 ⇒ rS2
lS1 ⇒ lS2
rS2 ⇒ rpIC
lS2 ⇒ lpIC
rpIC ⇒ raIC
rpIC ⇒ dAAC
lpIC ⇒ laIC
rpIC ⇒ dAAC
raIC ⇒ dAAC
laIC ⇒ dAAC
dAAC ⇒ rDLPFC
dAAC ⇒ lDLPFC
dAAC ⇒ mPFC
rS2 ⇒ rS1
lS2 ⇒ lS1
rDLPFC ⇒ dAAC
lDLPFC ⇒ dAAC
mPFC ⇒ dAAC
rDLPFC ⇒ lDLPFC
lDLPFC ⇒ rDLPFC
S1-r
S1-l
S2-r
S2-l
pIC-r
pIC-l
aIC-r
aIC-l
dAAC
DLPFC-r
DLPFC-l
mPFC
-10
10
[t]
20 40 60 80 [Hz]

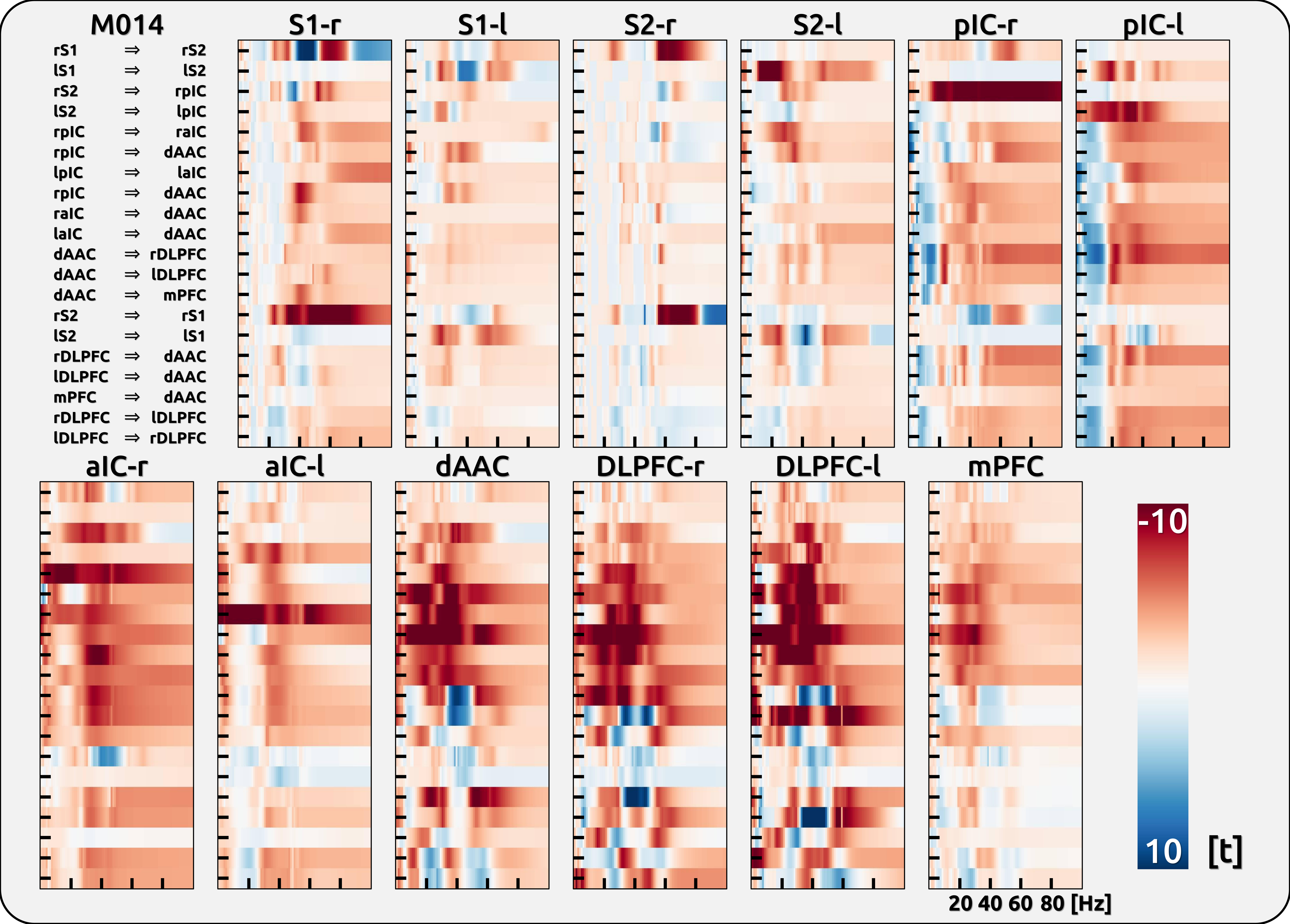

M014
S1-r
S1-l
S2-r
S2-l
pIC-r
pIC-l
rS1 ⇒ rS2
lS1 ⇒ lS2
rS2 ⇒ rpIC
lS2 ⇒ lpIC
rpIC ⇒ raIC
rpIC ⇒ dAAC
lpIC ⇒ laIC
rpIC ⇒ dAAC
raIC ⇒ dAAC
laIC ⇒ dAAC
dAAC ⇒ rDLPFC
dAAC ⇒ lDLPFC
dAAC ⇒ mPFC
rS2 ⇒ rS1
lS2 ⇒ lS1
rDLPFC ⇒ dAAC
lDLPFC ⇒ dAAC
mPFC ⇒ dAAC
rDLPFC ⇒ lDLPFC
lDLPFC ⇒ rDLPFC
aIC-r
aIC-l
dAAC
DLPFC-r
DLPFC-l
mPFC
-10
10
[t]
20 40 60 80 [Hz]

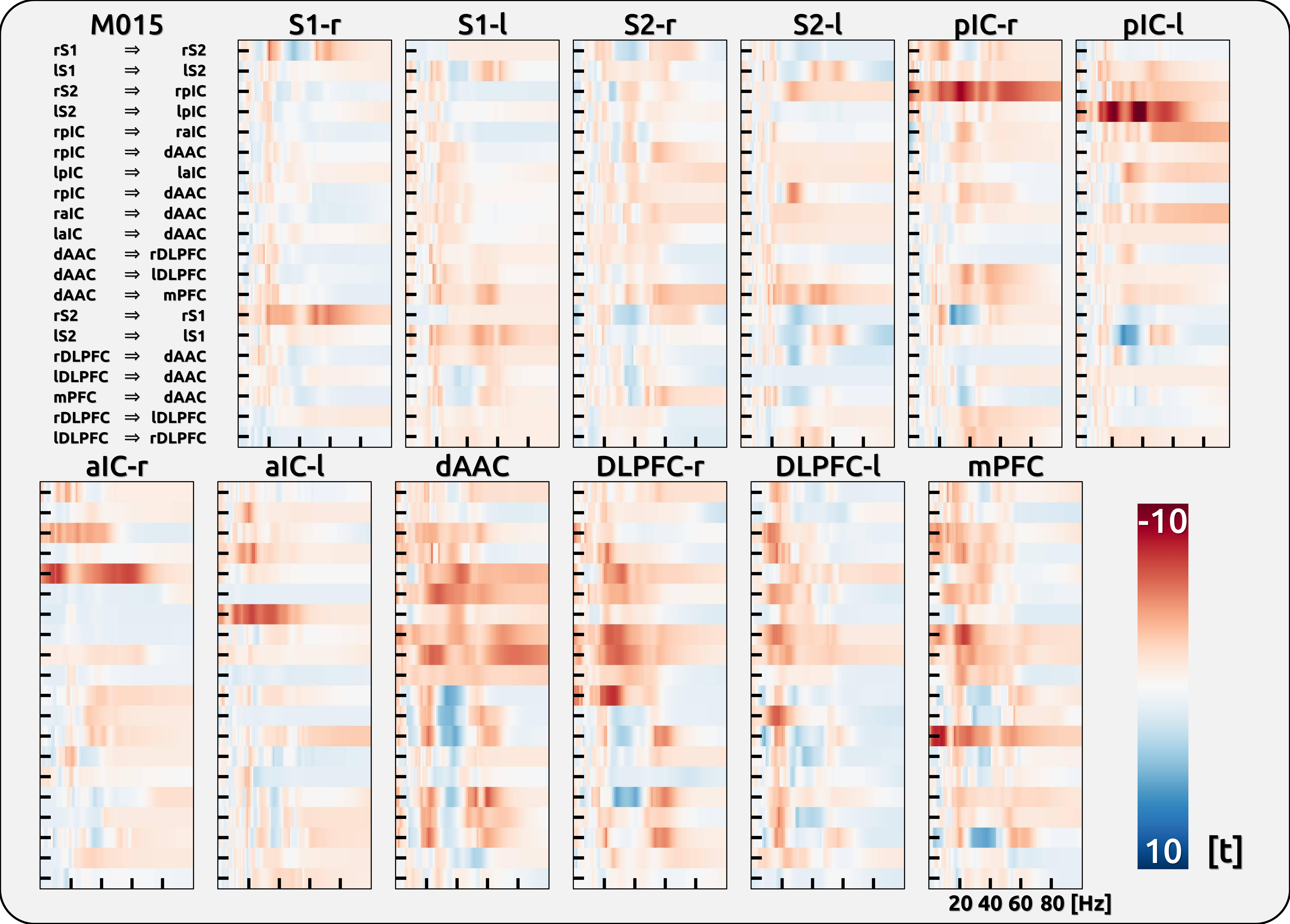

M015
S1-r
S1-l
S2-r
S2-l
pIC-r
pIC-l
rS1 ⇒ rS2
lS1 ⇒ lS2
rS2 ⇒ rpIC
lS2 ⇒ lpIC
rpIC ⇒ raIC
rpIC ⇒ dAAC
lpIC ⇒ laIC
rpIC ⇒ dAAC
raIC ⇒ dAAC
laIC ⇒ dAAC
dAAC ⇒ rDLPFC
dAAC ⇒ lDLPFC
dAAC ⇒ mPFC
rS2 ⇒ rS1
lS2 ⇒ lS1
rDLPFC ⇒ dAAC
lDLPFC ⇒ dAAC
mPFC ⇒ dAAC
rDLPFC ⇒ lDLPFC
lDLPFC ⇒ rDLPFC
aIC-r
aIC-l
dAAC
DLPFC-r
DLPFC-l
mPFC
-10
10
[t]
20 40 60 80 [Hz]

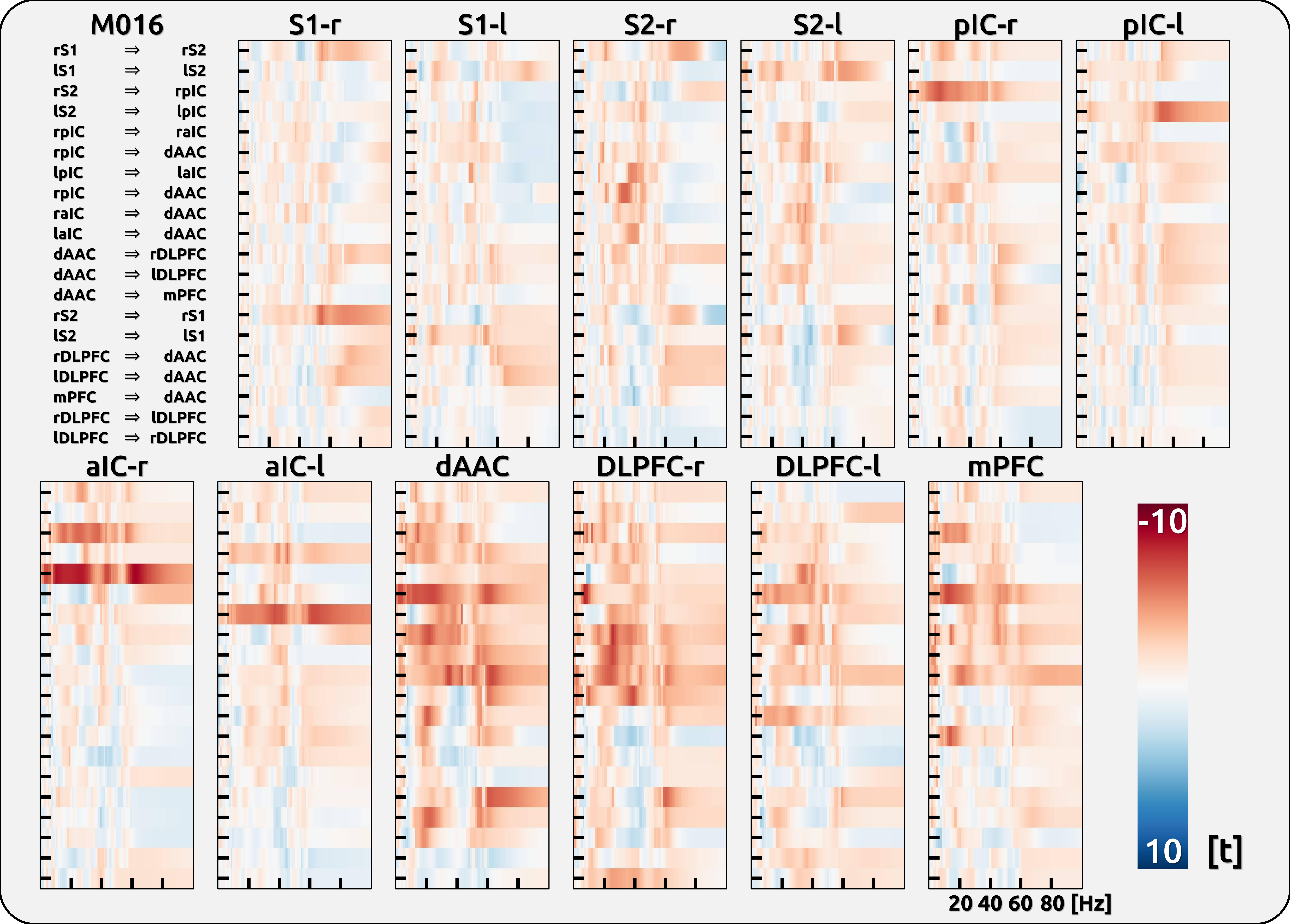

M016
S1-r
S1-l
S2-r
S2-l
pIC-r
pIC-l
rS1 ⇒ rS2
lS1 ⇒ lS2
rS2 ⇒ rpIC
lS2 ⇒ lpIC
rpIC ⇒ raIC
rpIC ⇒ dAAC
lpIC ⇒ laIC
rpIC ⇒ dAAC
raIC ⇒ dAAC
laIC ⇒ dAAC
dAAC ⇒ rDLPFC
dAAC ⇒ lDLPFC
dAAC ⇒ mPFC
rS2 ⇒ rS1
lS2 ⇒ lS1
rDLPFC ⇒ dAAC
lDLPFC ⇒ dAAC
mPFC ⇒ dAAC
rDLPFC ⇒ lDLPFC
lDLPFC ⇒ rDLPFC
aIC-r
aIC-l
dAAC
DLPFC-r
DLPFC-l
mPFC
-10
10
[t]
20 40 60 80 [Hz]

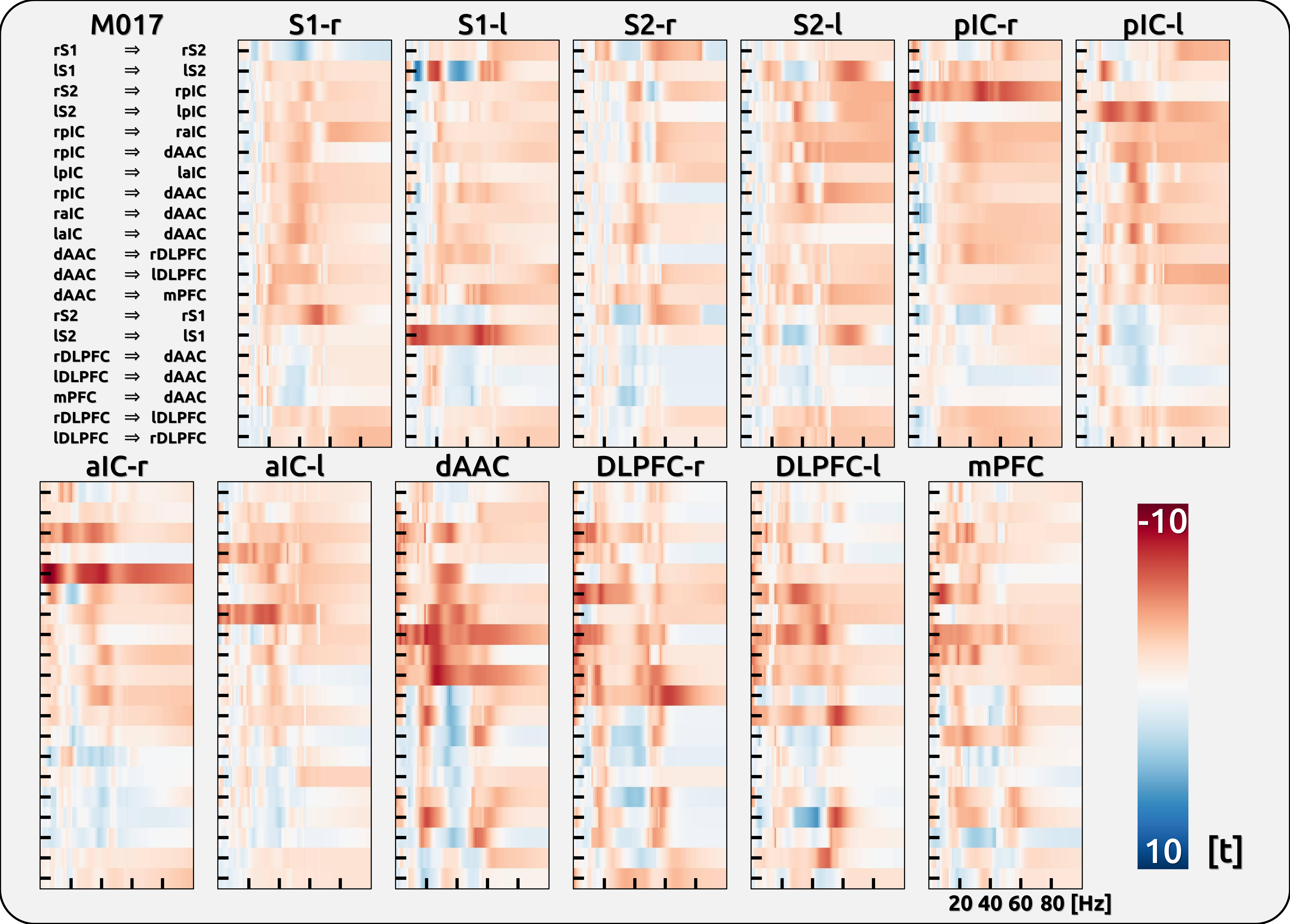

M017
S1-r
S1-l
S2-r
S2-l
pIC-r
pIC-l
rS1 ⇒ rS2
lS1 ⇒ lS2
rS2 ⇒ rpIC
lS2 ⇒ lpIC
rpIC ⇒ raIC
rpIC ⇒ dAAC
lpIC ⇒ laIC
rpIC ⇒ dAAC
raIC ⇒ dAAC
laIC ⇒ dAAC
dAAC ⇒ rDLPFC
dAAC ⇒ lDLPFC
dAAC ⇒ mPFC
rS2 ⇒ rS1
lS2 ⇒ lS1
rDLPFC ⇒ dAAC
lDLPFC ⇒ dAAC
mPFC ⇒ dAAC
rDLPFC ⇒ lDLPFC
lDLPFC ⇒ rDLPFC
aIC-r
aIC-l
dAAC
DLPFC-r
DLPFC-l
mPFC
-10
10
[t]
20 40 60 80 [Hz]

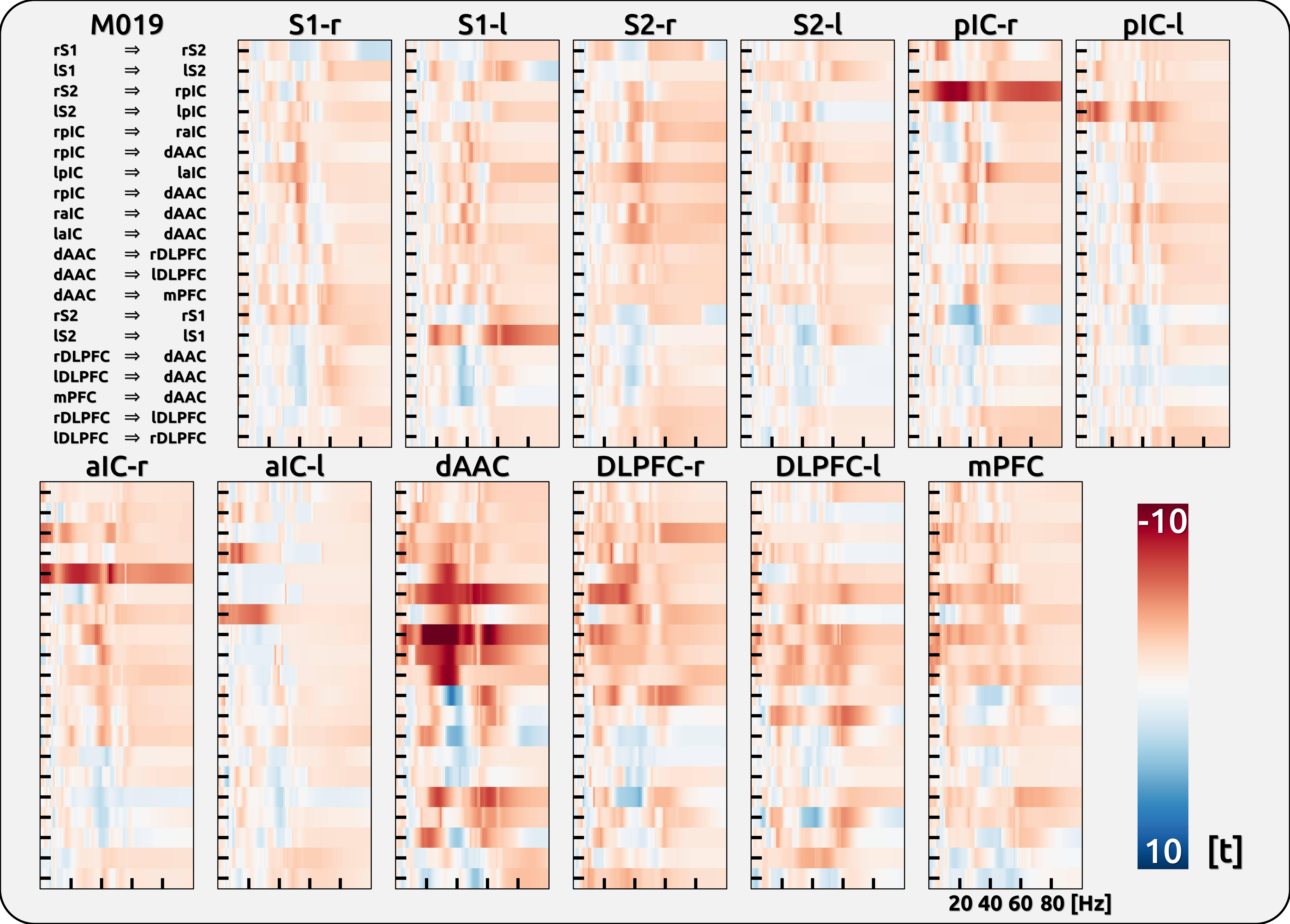

M019
S1-r
S1-l
S2-r
S2-l
pIC-r
pIC-l
rS1 ⇒ rS2
lS1 ⇒ lS2
rS2 ⇒ rpIC
lS2 ⇒ lpIC
rpIC ⇒ raIC
rpIC ⇒ dAAC
lpIC ⇒ laIC
rpIC ⇒ dAAC
raIC ⇒ dAAC
laIC ⇒ dAAC
dAAC ⇒ rDLPFC
dAAC ⇒ lDLPFC
dAAC ⇒ mPFC
rS2 ⇒ rS1
lS2 ⇒ lS1
rDLPFC ⇒ dAAC
lDLPFC ⇒ dAAC
mPFC ⇒ dAAC
rDLPFC ⇒ lDLPFC
lDLPFC ⇒ rDLPFC
aIC-r
aIC-l
dAAC
DLPFC-r
DLPFC-l
mPFC
-10
10
[t]
20 40 60 80 [Hz]

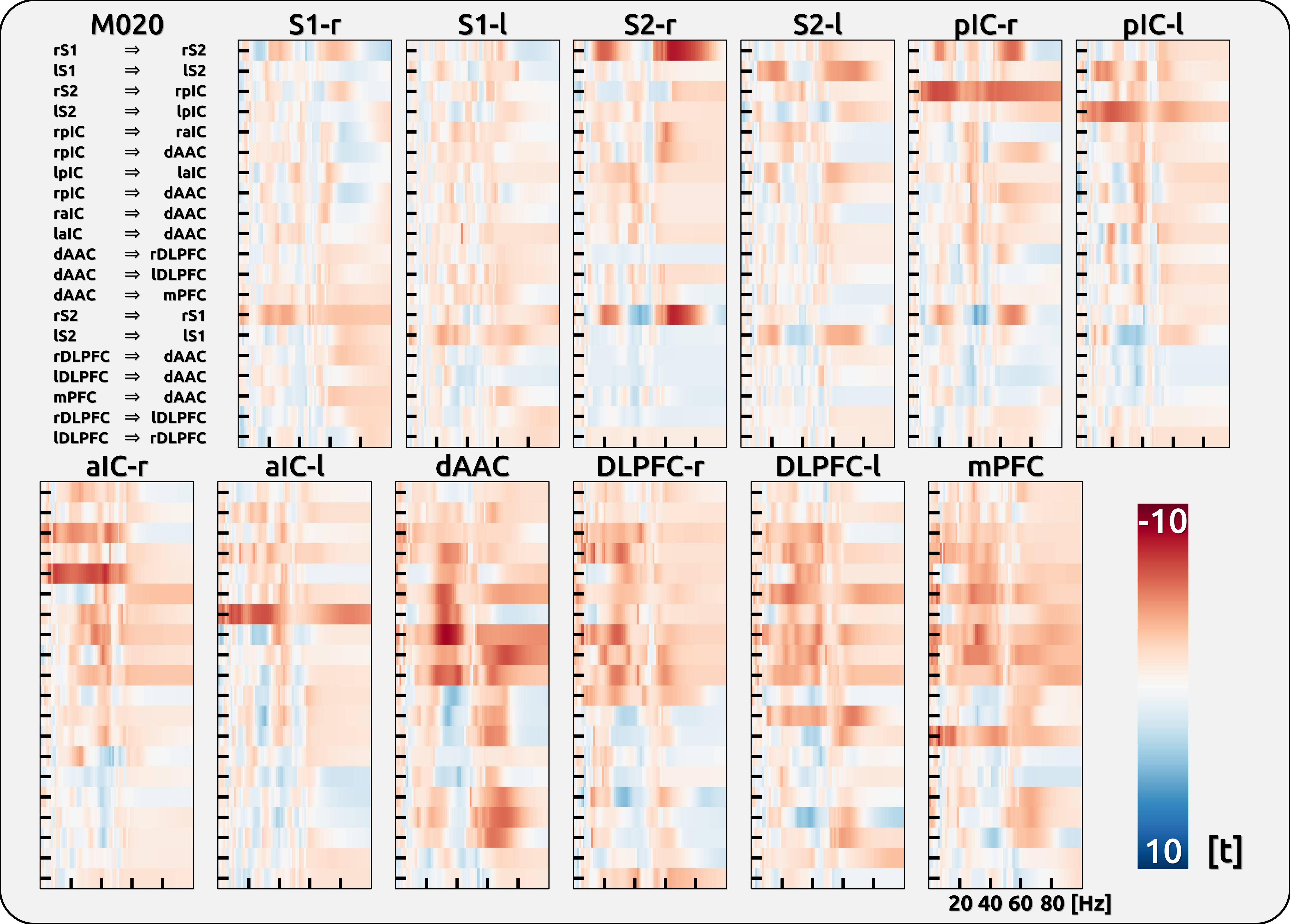
M020
rS1 ⇒ rS2
lS1 ⇒ lS2
rS2 ⇒ rpIC
lS2 ⇒ lpIC
rpIC ⇒ raIC
rpIC ⇒ dAAC
lpIC ⇒ laIC
rpIC ⇒ dAAC
raIC ⇒ dAAC
laIC ⇒ dAAC
dAAC ⇒ rDLPFC
dAAC ⇒ lDLPFC
dAAC ⇒ mPFC
rS2 ⇒ rS1
lS2 ⇒ lS1
rDLPFC ⇒ dAAC
lDLPFC ⇒ dAAC
mPFC ⇒ dAAC
rDLPFC ⇒ lDLPFC
lDLPFC ⇒ rDLPFC
S1-r
S1-l
S2-r
S2-l
pIC-r
pIC-l
aIC-r
aIC-l
dAAC
DLPFC-r
DLPFC-l
mPFC
-10
10
[t]
20 40 60 80 [Hz]

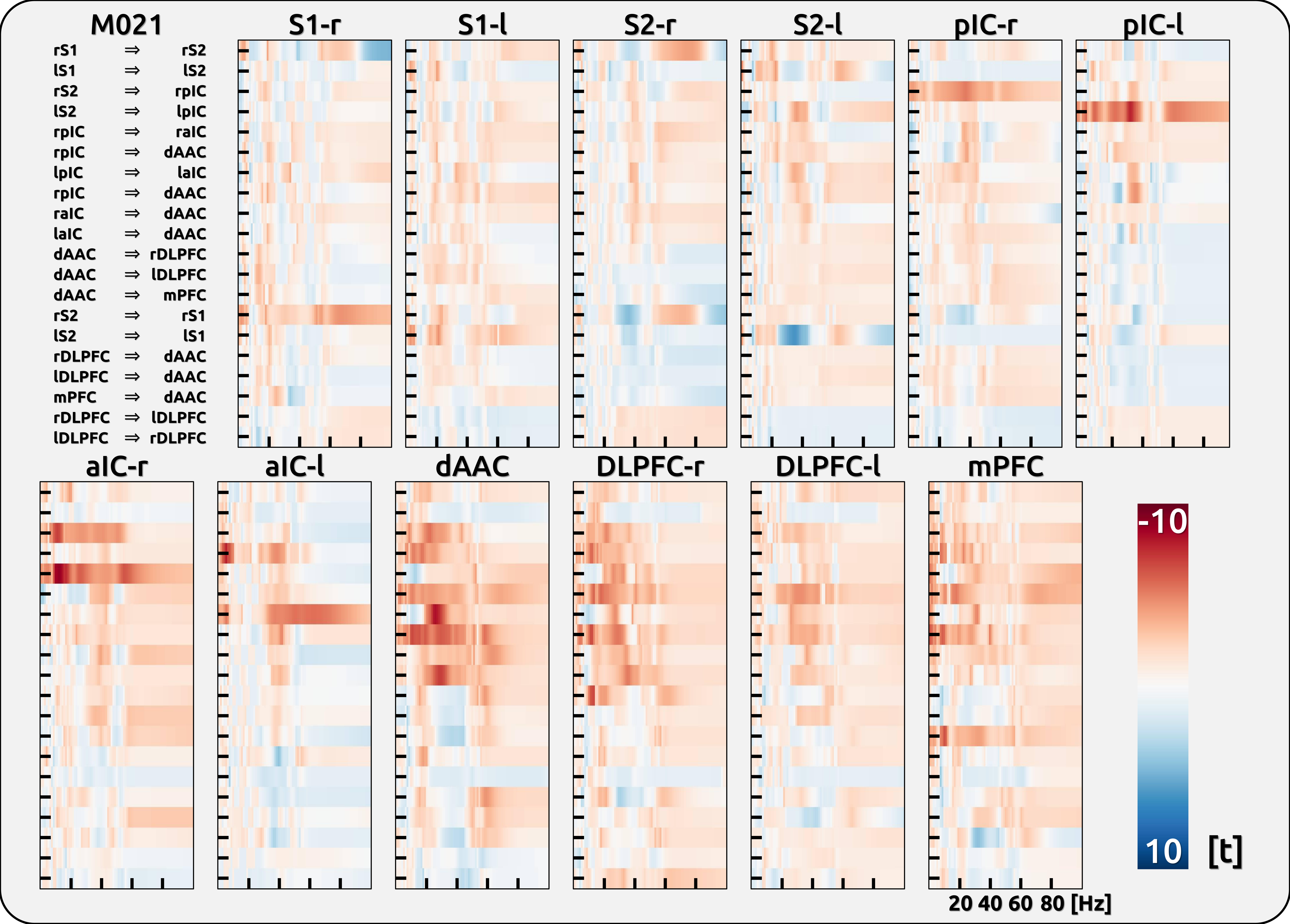
M021
S1-r
S1-l
S2-r
S2-l
pIC-r
pIC-l
rS1 ⇒ rS2
lS1 ⇒ lS2
rS2 ⇒ rpIC
lS2 ⇒ lpIC
rpIC ⇒ raIC
rpIC ⇒ dAAC
lpIC ⇒ laIC
rpIC ⇒ dAAC
raIC ⇒ dAAC
laIC ⇒ dAAC
dAAC ⇒ rDLPFC
dAAC ⇒ lDLPFC
dAAC ⇒ mPFC
rS2 ⇒ rS1
lS2 ⇒ lS1
rDLPFC ⇒ dAAC
lDLPFC ⇒ dAAC
mPFC ⇒ dAAC
rDLPFC ⇒ lDLPFC
lDLPFC ⇒ rDLPFC
aIC-r
aIC-l
dAAC
DLPFC-r
DLPFC-l
mPFC
-10
10
[t]
20 40 60 80 [Hz]

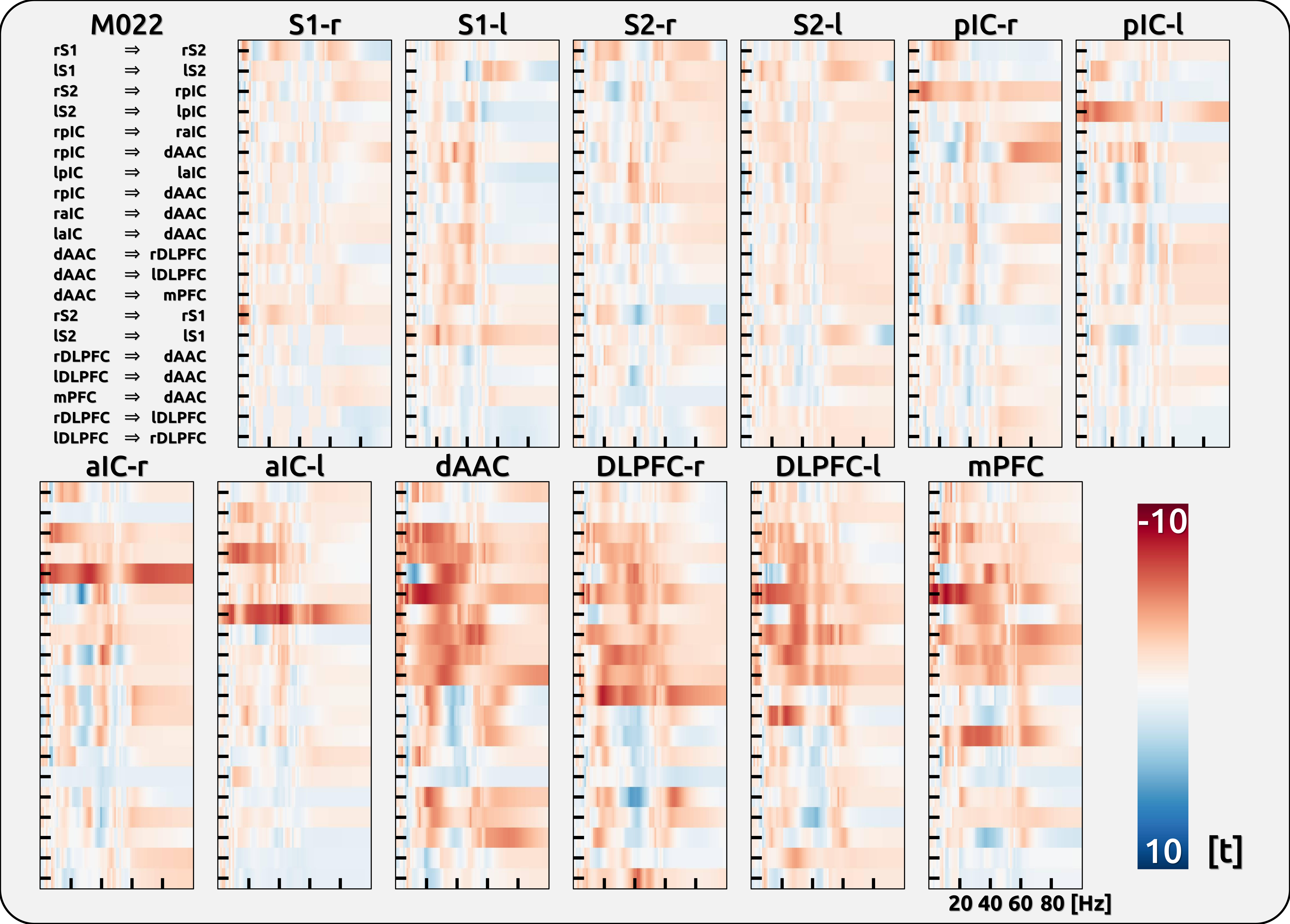

M022
rS1 ⇒ rS2
lS1 ⇒ lS2
rS2 ⇒ rpIC
lS2 ⇒ lpIC
rpIC ⇒ raIC
rpIC ⇒ dAAC
lpIC ⇒ laIC
rpIC ⇒ dAAC
raIC ⇒ dAAC
laIC ⇒ dAAC
dAAC ⇒ rDLPFC
dAAC ⇒ lDLPFC
dAAC ⇒ mPFC
rS2 ⇒ rS1
lS2 ⇒ lS1
rDLPFC ⇒ dAAC
lDLPFC ⇒ dAAC
mPFC ⇒ dAAC
rDLPFC ⇒ lDLPFC
lDLPFC ⇒ rDLPFC
S1-r
S1-l
S2-r
S2-l
pIC-r
pIC-l
aIC-r
aIC-l
dAAC
DLPFC-r
DLPFC-l
mPFC
-10
10
[t]
20 40 60 80 [Hz]

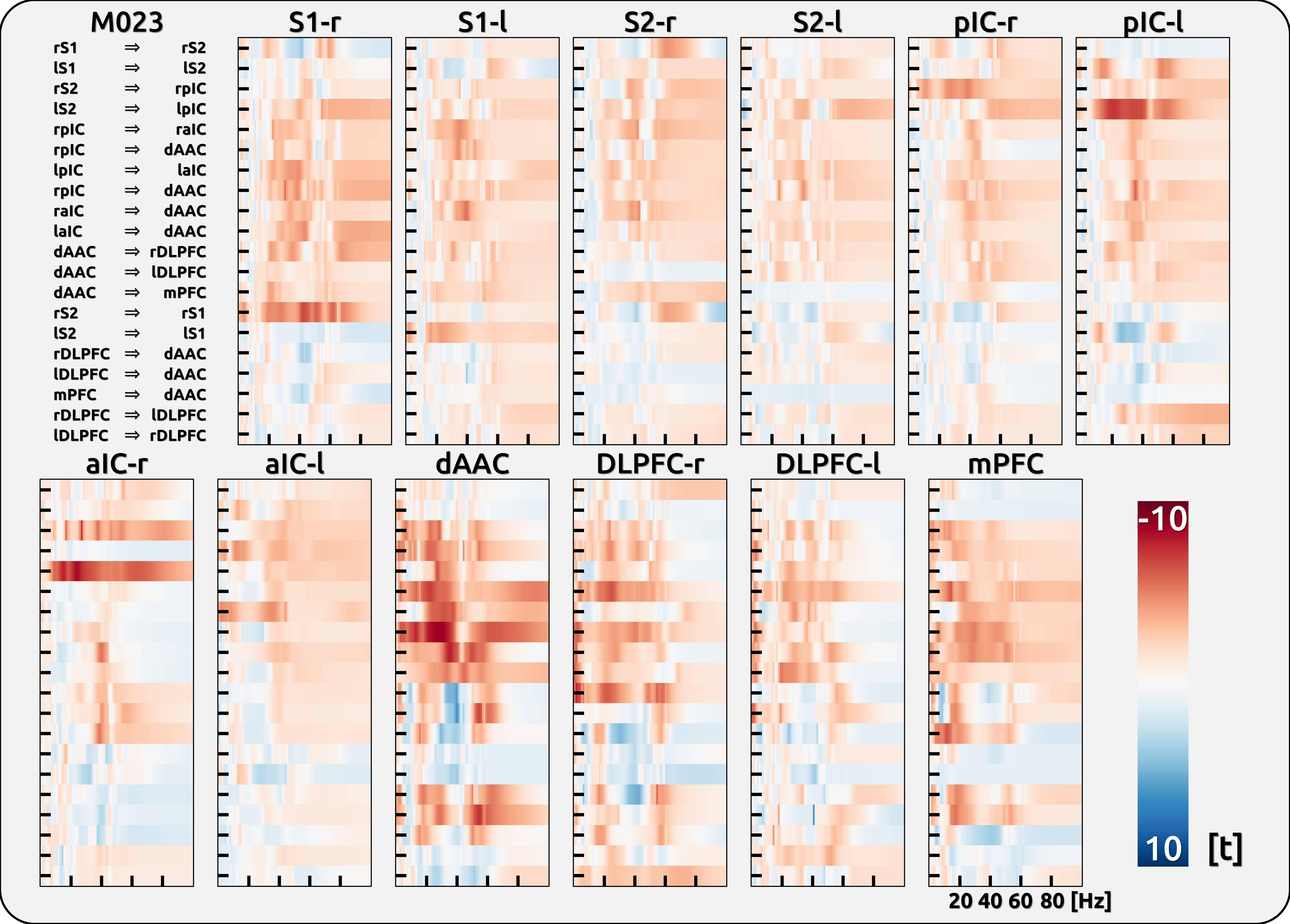
M023
rS1 ⇒ rS2
lS1 ⇒ lS2
rS2 ⇒ rpIC
lS2 ⇒ lpIC
rpIC ⇒ raIC
rpIC ⇒ dAAC
lpIC ⇒ laIC
rpIC ⇒ dAAC
raIC ⇒ dAAC
laIC ⇒ dAAC
dAAC ⇒ rDLPFC
dAAC ⇒ lDLPFC
dAAC ⇒ mPFC
rS2 ⇒ rS1
lS2 ⇒ lS1
rDLPFC ⇒ dAAC
lDLPFC ⇒ dAAC
mPFC ⇒ dAAC
rDLPFC ⇒ lDLPFC
lDLPFC ⇒ rDLPFC
S1-r
S1-l
S2-r
S2-l
pIC-r
pIC-l
aIC-r
aIC-l
dAAC
DLPFC-r
DLPFC-l
mPFC
-10
10
[t]
20 40 60 80 [Hz]
